# Coding of social novelty in the hippocampal CA2 region and its disruption and rescue in a mouse model of schizophrenia

**DOI:** 10.1101/833723

**Authors:** Macayla L. Donegan, Fabio Stefanini, Torcato Meira, Joshua A. Gordon, Stefano Fusi, Steven A. Siegelbaum

## Abstract

The hippocampal CA2 region is essential for social memory and has been implicated in neuropsychiatric disorders. However, little is known about how CA2 neural activity encodes social interactions and how this coding is altered in disease. We recorded from CA2 pyramidal neurons as mice engaged in social interactions and found that while CA2 failed to stably represent spatial location, CA2 activity encoded contextual changes and novel social stimuli. In the *Df(16)A^+/-^* mouse model of the human 22q11.2 microdeletion, a major schizophrenia risk factor, CA2 activity showed a surprising increase in spatial coding while failing to encode social novelty, consistent with the social memory deficit in these mice. Previous work has shown that CA2 pyramidal neurons are hyperpolarized in *Df(16)A^+/-^* mice, likely as a result of upregulation of TREK-1 K^+^ current. We found that administration of a TREK-1 antagonist rescued the social memory deficits and restored normal CA2 coding properties in *Df(16)A^+/-^* mice, supporting a crucial role for CA2 in the encoding of novel social stimuli and social dysfunction.

## Introduction

Social memory, the ability of an animal to recognize and remember a previously encountered conspecific, is indispensable for a wide range of social behaviors^1^. Deficits in social memory and social behavioral changes are commonly associated with neuropsychiatric disease^2^. Lesion studies in both humans^3^ and rodents^4^ indicate that the hippocampus, a brain region well-known to be important for several forms of declarative memory^5^, is also necessary for encoding social memory. Although the ability of hippocampal neural firing to represent spatial, contextual and semantic information that may contribute to memory encoding has been well established^6–9^, how hippocampus encodes and represents social information in behaviorally relevant contexts is less well understood.

Recent studies indicate that the long overlooked hippocampal CA2 subregion is a critical component of the circuit necessary for encoding social information into declarative memory. Both short and long-term silencing of dorsal CA2 prevents social memory formation, consolidation and recall^10–12^. Although dorsal CA2 provides strong input to dorsal CA1, recently our laboratory found that social memory depends on the CA2 projections to ventral CA1^10^, an area that is also required for social memory and that can encode social engrams^13, 14^. However, at present, it is unclear as to whether and how dorsal CA2 itself encodes social information that could be relevant for social memory.

Previous studies have found that CA2 spatial firing properties clearly differ from those of neighboring dorsal CA1 and CA3 regions. Thus, CA2 place fields have lower spatial information than those in CA1 or CA3^15–18^. CA2 place fields are spatially unstable in the same environment over time^15^, showing clear differences with the more stable and spatially precise CA1 place cell firing. CA2 activity is also more sensitive to contextual change than CA1 and CA3^19^. Of interest, CA2 place fields globally remap in the presence of novel objects or of familiar or novel social stimuli^16^, although whether CA2 firing contains specific social information that is relevant to social memory remains unkown.

The role of CA2 in social memory is of particular clinical relevance as postmortem hippocampal tissue from individuals with schizophrenia or bipolar disorder reveal a 30% decrease in the number of parvalbumin positive interneurons selectively in CA2, with no changes in other hippocampal regions^20, 21^. A similar CA2-selective loss of PV+ interneurons is observed in the *Df(16)A^+/-^* mouse model of the human 22q11.2 microdeletion^22^, which confers a 30-fold increase in the risk of developing schizophrenia^23^. Although the decrease in inhibition might be expected to enhance CA2 pyramidal neuron (PN) activity and thus enhance social memory, these mice actually have a profound deficit in social memory^22^. This behavioral deficit may reflect the fact that, in addition to the decreased inhibition, CA2 PNs become hyperpolarized in these mice, likely as a result of the upregulation of the TREK-1 two-pore K+ channel, which is normally highly enriched in CA2^22^. It remains an open question as to whether and how the opposing actions of decreased CA2 inhibition and hyperpolarization of CA2 principal cells affect their *in vivo* firing properties. Additionally, the role of TREK-1 upregulation in the social memory deficits seen in the *Df(16)A^+/-^* mice has not been explored at either the behavioral or electrophysiological level.

Here we examined CA2 firing properties using extracellular recordings from dorsal CA2 pyramidal neurons in freely behaving animals during spatial exploration and social interactions. We found that CA2 neurons failed to encode location but reliably reported social novelty at both the single cell and population level in wild-type mice. Moreover, social novelty coding was impaired in the *Df(16)A^+/-^* mice, providing a potential explanation for their social memory deficit^22^. Surprisingly, CA2 neurons in the *Df(16)A^+/-^* mice showed enhanced spatial coding properties, suggesting that neuropsychiatric disease can transform the major cognitive function of a given brain region. Finally, we found that suppression of TREK-1 in the disease model mice rescued both normal social memory behavior and the normal social and spatial coding preferences of CA2 neurons, suggesting that CA2 neurons may provide a novel drug target for treating the negative social symptoms of schizophrenia.

## Results

### CA2 spatial firing was unstable during a three-chamber social interaction task

We characterized CA2 firing as mice investigated various social and non-social stimuli by performing extracellular electrophysiological recordings from dorsal CA2 and CA1 pyramidal neurons during a three-chamber social interaction task (Figure 1a). In the task a mouse was exposed to a three-chamber arena in five sequential 10 minute sessions in which: 1) the chambers were void of all objects (empty chamber session); 2) the chambers contained two identical wire cup cages as novel objects in the two end chambers (objects session); 3) two familiar littermates (L1 and L2) were placed one each in the wire cup cages (familiar social session 1); 4) one littermate was replaced with a novel mouse (novel social session); and 5) the novel mouse was replaced by the original littermate (familiar social session 2).

**Figure 1.**
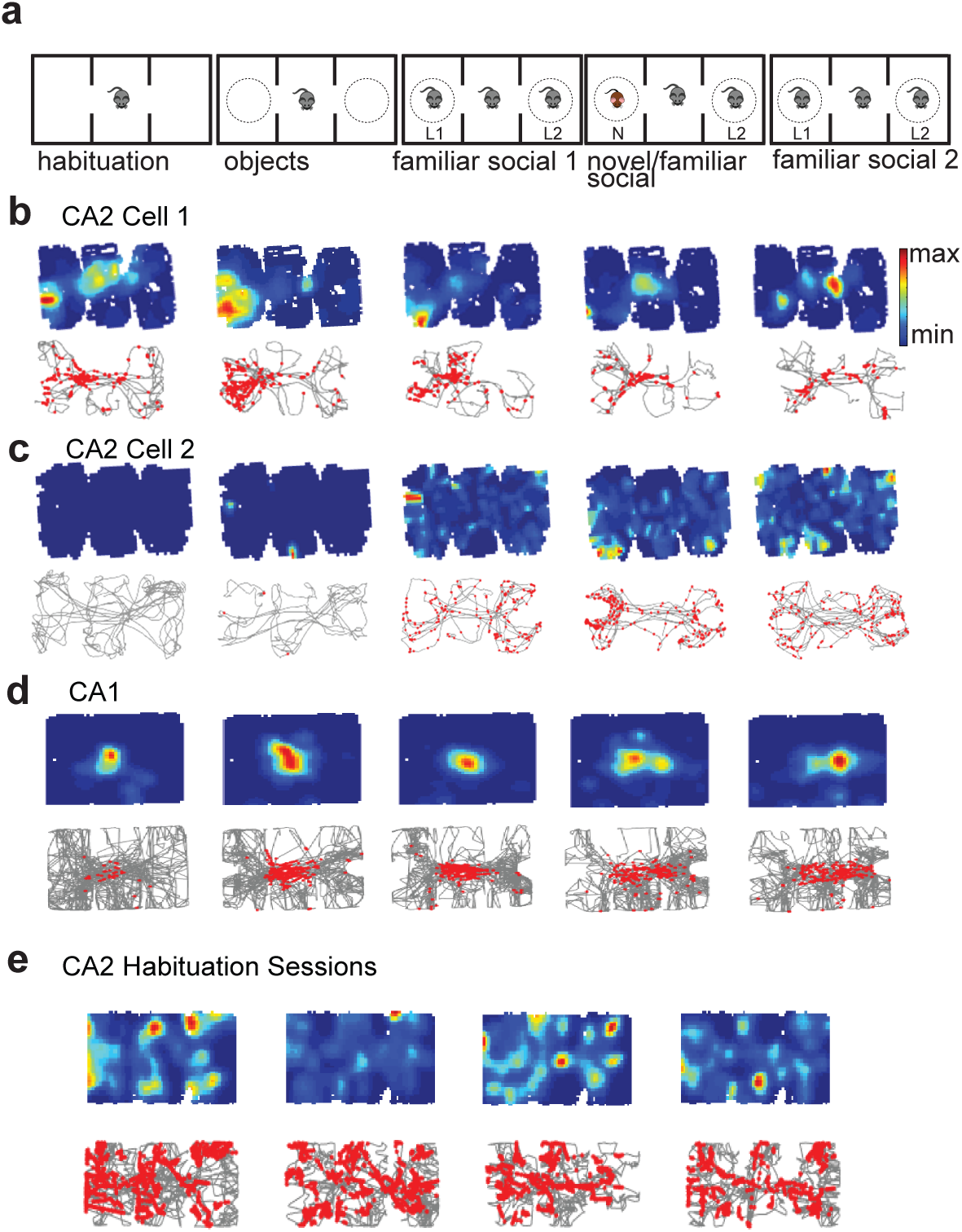
Hippocampal firing in the three-chamber interaction task. (a) The three-chamber interaction task. Mice explored in five sequential 10-min sessions the three chamber environment: 1) empty arena (empty session), 2) two identical novel objects (empty wire cup cage) placed in the two side chambers (objects session), 3) two familiar littermates (L1 and L2) placed one in each cup (familiar social session 1; fam1), 4) a novel mouse (N) present in one cup and the remaining littermate present in the other cup (novel session), 5) return to the two original familiar mice (familiar session 2; fam2). (b) Example CA2 neuron place cell heat maps (top). Bottom, single spikes (red dots) on top of trajectory trace (gray). Maximum firing rates in sessions 1-5 were: 7 Hz, 7 Hz, 9 Hz, 9 Hz, 5 Hz, respectively. (c) Example CA2 neuron that was nearly silent in non-social sessions 1 and 2 and that became active in social sessions 3-5. Maximum firing rates in sessions 1-5 were: 1 Hz, 2 Hz, 15 Hz, 22 Hz, 10 Hz. (d) Example CA1 cell showing stable place fields throughout all sessions. Maximum firing rates in sessions 1-5 were: 1 Hz, 5 Hz, 7 Hz, 4 Hz, 4 Hz. (e) Example firing of a CA2 cell during four 10-min successive sessions of a 40 min habituation period to the empty chambers. Maximum firing rates in sessions 1-4 were: 19 Hz, 34 Hz, 13 Hz, 19 Hz.

We found that CA2 neurons showed only weak spatially selective firing as an animal explored the three chambers in the five sessions (Figure 1b,c), consistent with previous reports under different behavioral conditions^15, 16, 24^. CA2 place fields were large and diffuse, with a typical neuron exhibiting multiple firing fields that were less spatially selective than CA1 fields in the same environment (Figure 1b-d). Analysis of mean data from 192 CA2 neurons from 6 animals and 87 CA1 neurons from 3 animals revealed that CA2 PNs have more place fields per cell (CA2 = 2.61 ± 0.12 fields; CA1 = 2.0 ± 0.15 fields; p=0.02, paired t-test), larger place fields (CA2=107.3 ± 10.1 pixels; CA1=62.94 ± 5.96 pixels; p=0.04, paired t-test), and lower spatial information scores than CA1 (CA2 = 0.42 ± 0.02 bits/spike; CA1 = 0.60 ± 0.09 bits/spike; p < 0.001, paired t-test). This is in agreement with previous studies comparing CA2 firing properties to those of CA1 in an open field^15, 24^. The number and size of CA2 place fields, along with the amount of spatial information, did not vary from session to session in the three-chamber task (Supplemental Figure 1).

In addition to the decreased spatial information compared to CA1, CA2 place fields were less spatially stable across the different sessions of the three-chamber task in comparison to CA1. This was evident in both individual cell firing plots (Figure 1b-d), and in measurements of Pearson’s correlation values (r) of place fields between different sessions (Figure 2a, c). Of interest, CA2 spatial firing was significantly more stable during four identical 10-min sessions throughout a 40-minute-long habituation period to the empty three-chamber environment carried out on the day prior to the three-chamber task (Figures 1e and 2b,c), suggesting that the spatial instability across the different sessions of the three-chamber task is related to alterations in the content of the chambers (Figures 1f,2b).

**Figure 2.**
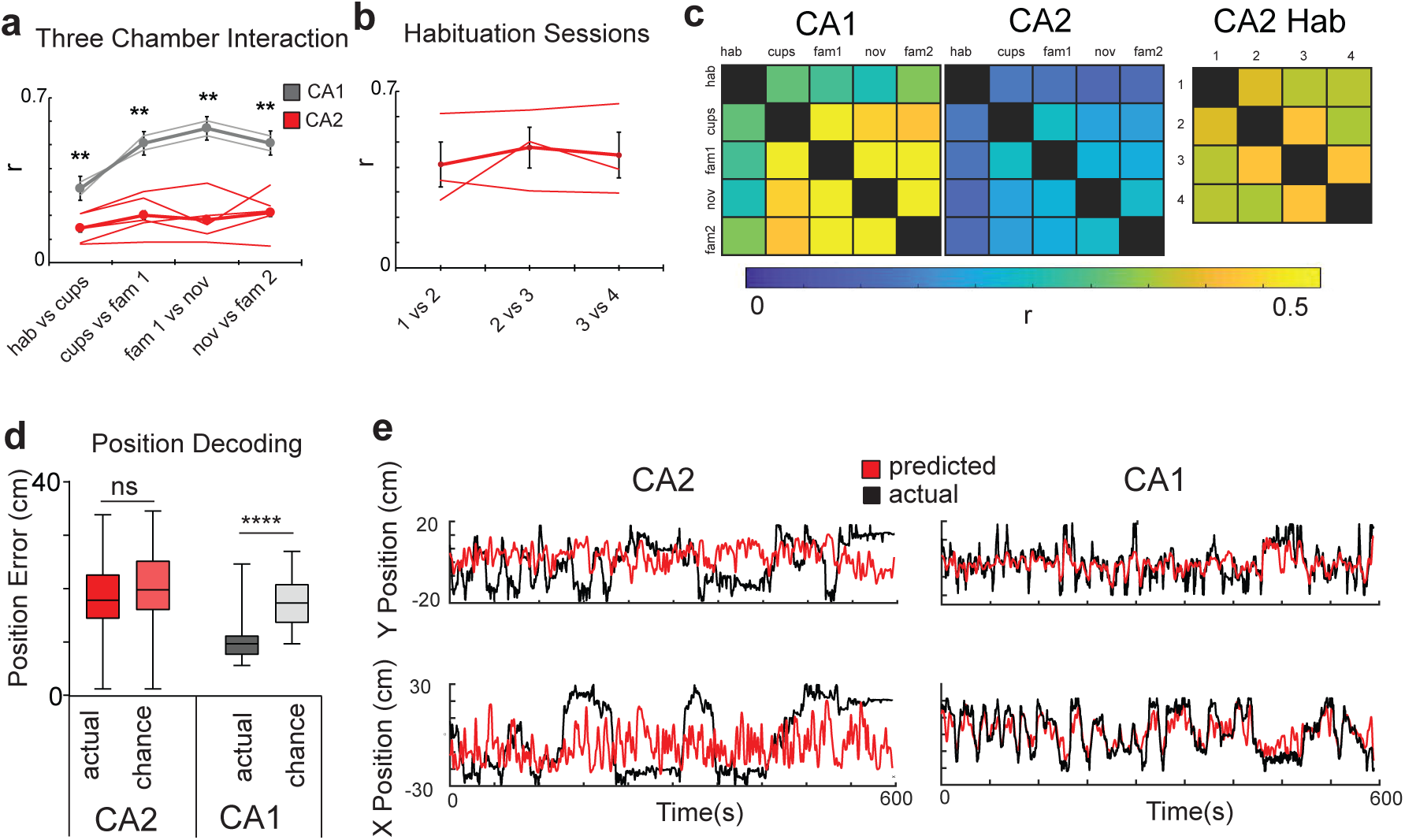
CA2 spatial firing is unstable and fails to decode position. (a) Plot of Pearson’s correlation (r) between place field maps in successive pairs of the five three-chamber task sessions for CA2 and CA1 neurons. Thin traces show results from individual animals and thick traces show means (n=192 CA2 neurons from 6 animals; n=87 CA1 neurons from 3 animals). Error bars show SEM. CA2 firing was less stable than CA1 firing (paired t-tests with Bonferroni correction for multiple comparisons; p = 0.02, 0.006, 0.003, 0.009). (b) CA2 place field correlations between pairs of the four 10-min habituation sessions. (c) Left, Color-coded plots of mean spatial correlations between each pair of sessions averaged over all CA2 and CA1 neurons in three-chamber task. Right, CA2 neuron correlations for pairs of sessions during the 40-min habituation period. (d) Mean error of position with support vector machine (SVM) decoding based on CA2 and CA1 population spatial firing data in three-chamber task compared to chance performance. CA1 decoding performed above chance (p<.0001, Wilcoxon rank-sum test; n=87 neurons from 3 animals) whereas CA2 did not (p>0.05; n=192 neurons from 6 animals).(e) Example of SVM position decoding with a linear kernel from CA2 and CA1 neuron firing from individual mice. Actual X-Y position trajectory (black traces) and predicted location (red traces) decoded from CA2 and CA1 firing (smoothed for visualization). Location plotted relative to center of the three-chamber environment.

It has been suggested that CA2 spatial firing becomes more stable after the addition of a social stimulus^16^, which would imply that spatial firing patterns would be more similar between sessions with the same social stimuli. However, we found that the spatial correlation between the two familiar mice sessions (sessions 3 versus 5; r = 0.21 ± 0.22), which contained the same social stimuli at the same locations, was no greater than the spatial correlations between the object and familiar social sessions (sessions 2 versus 3/5; r = 0.23 ± 0.24), or the novel and familiar social sessions (sessions 3 versus 4; r = 0.25 ± 0.27; p>0.05 in all comparisons, paired t-tests). These results indicate that the stability of spatial information that CA2 encodes was not enhanced by contextual information (Figure 2a,c).

### CA2 population activity encoded contextual but not spatial information in the three-chamber task

Several studies have shown that populations of neurons may accurately encode aspects of an environment even if individual neuron firing properties do not. Thus, place fields in dentate gyrus^25^ and ventral CA1^26^, which have lower spatial information content compared to dorsal CA1 neurons, can nonetheless encode position at the population level as accurately as dorsal CA1. To examine whether this was the case for dorsal CA2 pyramidal neurons, we used a machine learning approach. A set of support vector machines (SVM)^27–29^ using a linear kernel were trained to decode the position of an animal as it explored the three chambers based on CA1 or CA2 population activity. Whereas the decoder based on CA1 activity accurately predicted an animal’s location in all sessions of the three-chamber task, the decoder based on CA2 activity failed to predict spatial location above chance levels during any of the sessions (Figure 2c; Supplemental Figure 2). CA2 also failed to decode position in any of the four 10-min sessions of the 40 min period of habituation to the empty chambers, indicating that CA2 also provided weak spatial representations of a constant, empty environment (Supplemental Figure 2).

Next we examined whether the sensitivity of CA2 firing to contextual change^19^ was sufficient to decode the changes in environmental content in the five sessions of the three-chamber task, and whether there were any differences in decoding ability of CA2 compared to CA1. Indeed, when we trained a linear decoder using CA2 population activity to determine in which session an animal was engaged, the decoder performed significantly better than chance. Moreover, the CA2-based decoder performed significantly better than the CA1-based decoder in identifying task session (Figure 3a,b).

**Figure 3.**
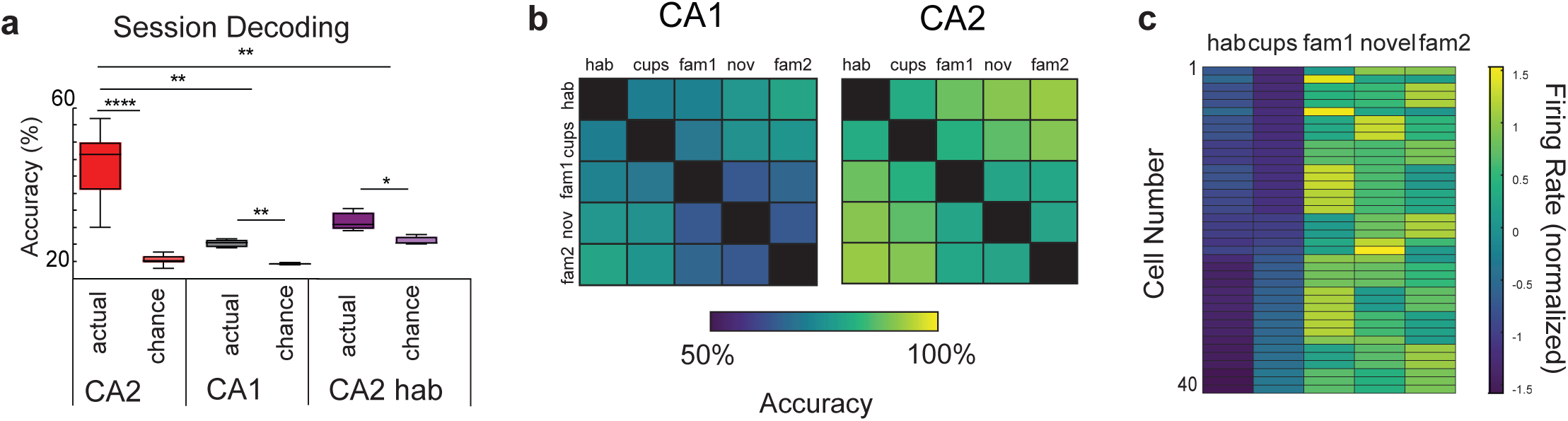
CA2 encodes contextual changes and the presence of social stimuli. (a) SVM decoder performance for identifying the particular session in which a mouse was engaged in the three-chamber task or during the four 10-min sessions of empty chamber habituation (hab). Decoder trained on either CA2 (dark red) or CA1 (dark grey) firing performed significantly better than chance (lighter shaded bars): CA2, p < 0.001 (n=192 neurons from 6 animals); CA1, p < 0.01 (n=87 neurons from 3yy animals). CA2 three-chamber session decoding accuracy was significantly greater than CA1 (p < 0.01, Wilcoxon rank-sum test). Decoder trained on CA2 activity during four 10-min sessions of habituation period predicted habituation session slightly above chance (p=0.03, n=94 neurons from 4 animals). Accuracy of habituation session decoding was significantly lower than that for three-chamber session decoding (p<0.01). The ratio of performance over chance accuracy was significantly higher in the three-chamber sessions (2.1 ± 0.09) compared to habituation sessions (1.2 ± 0.06; p=0.002). (b) CA1 and CA2 color-coded decoding accuracy for all possible session pairs in three-chamber task. (c) 40/192 CA2 PNs significantly increased their firing rate in the presence of social stimuli (difference in normalized firing rate > 2 standard deviations from the non-social sessions to the social sessions, see Supplemental Figure 2 for more information).

The ability of CA2 activity to distinguish among the different sessions could result from a sensitivity of neuronal firing to the differences in environmental content (context) between successive sessions (e.g. presence of non-social or social cues). Alternatively, it could reflect a sensitivity of CA2 firing to the passage of time independent of environmental content^15^. To distinguish between these possibilities, we trained a decoder on CA2 activity in the 4 sessions of the 40 min habituation period, where there was no change in content but where the passage of time was similar to that in the three-chamber task. Although the decoder was able to distinguish among the four habituation sessions slightly above chance level (p=0.03), decoder accuracy was significantly below that observed for the three-chamber task (p=0.01; Figure 3a), with the ratio of performance accuracy to chance accuracy in the three-chamber task (2.1 ± 0.09) significantly larger than in the habituation sessions (1.2 ± 0.06; p=0.002, t-test). We thus conclude that CA2 contains significant information about environmental content in addition to any information about the passage of time. This conclusion is further supported by our finding that CA2 place fields were more stable in the 40 min habituation session compared to the three-chamber task (Figure 2 a,b) and by additional findings based on social coding properties presented below.

### A subset of CA2 cells increased their firing rate in the presence of a social stimulus

We next explored whether CA2 firing was sensitive to the presence of another mouse (a social stimulus). The mean z-scored firing rate of CA2 neurons differed significantly among the non-social and social sessions of the three-chamber task (ANOVA; p=0.0004), whereas CA1 firing remained relatively constant (ANOVA p=0.29). Moreover, 40/192 (∼20%) of individual CA2 neurons significantly increased their mean z-scored firing rate (>2) during the social sessions (3-5) compared to the non-social sessions (1 and 2) (Figure 3c). In addition, 12 of the 40 neurons were initially silent (or nearly so) in the two preceding non-social sessions, with an initial firing rate in bottom 5% of the population (<0.007 Hz; Supplemental Figure 3). The increase in activity did not reflect random shifts in firing as only 3/192 (<2%) of CA2 cells were significantly more active during the non-social sessions than the social sessions, and none fell silent during the social sessions. In contrast to the enhanced social firing of CA2 neurons, only 2/87 CA1 cells showed a significant increase in firing rate (z-score >2) in the social sessions compared to the preceding non-social sessions, similar to the 3/87 fraction of CA1 cells that fired significantly more during the non-social sessions.

Previous studies have found that CA2 firing responds to both novel objects and social stimuli^16^ whereas silencing of CA2 impairs social but not object memory^11^. These separate results suggest that objects and social stimuli may differentially engage CA2 firing. We thus examined whether the population response of CA2 firing to novel objects differed from its firing to social stimuli by comparing CA2 population firing in the session containing the wire cup cages (novel objects) to the sessions containing the social stimuli. For each CA2 neuron, we determined the change in z-scored firing rate between a given session (objects or social stimuli) and its initial firing rate in the empty arena (session 1) and then expressed this information as population firing rate vectors to the objects or social stimuli. There was a highly significant difference in the CA2 population vectors for the firing rate changes in the object session compared to the social sessions (p= 5.8 × 10^-37^; Wilcoxon rank-sums test), indicating that the population of CA2 neurons was indeed differentially responsive to the addition of a social stimulus compared to an inanimate object (Supplemental Figure 3).

### CA2 encodes social novelty

To determine whether CA2 may encode specific social information that could contribute to social memory, we focused on CA2 firing while an animal was exploring within a body’s length (7 cm) of the cups (the interaction zone), either with or without a caged mouse present (Figure 4a). To limit potential spatial firing contributions, we compared CA2 firing among different sessions within a single interaction zone, defined by the cup that contained the novel mouse in session 4. We found that 77/192 CA2 PNs showed a significant (>2 SD) increase in firing during interactions with a novel mouse compared to a familiar mouse, as seen in both color-coded plots of neuronal firing-rate versus time (Figure 4b) and in scatter plots of mean firing rates (Figure 4c). Some CA2 PNs maintained an increase in firing around the novel animal throughout the 10 min period of a given social session whereas other neurons increased their firing only transiently, during the initial encounters with the novel animal (Figure 4b). As a result of the contributions of such neurons, mean CA2 population z-scored firing rate when an animal was exploring within the interaction zone around a novel animal (0.83 ± 0.06) was significantly greater than the firing rate around the familiar littermate in the flanking sessions (−0.26 ± 0.04; Mann Whitney U<0.0001) (Figure 4b,c). Moreover, the population firing rate vector around the novel animal also differed significantly from the firing rate vector around the familiar animal (averaged from sessions 3 and 5; p<0.0001, Wilcoxon rank-sum test).

**Figure 4.**
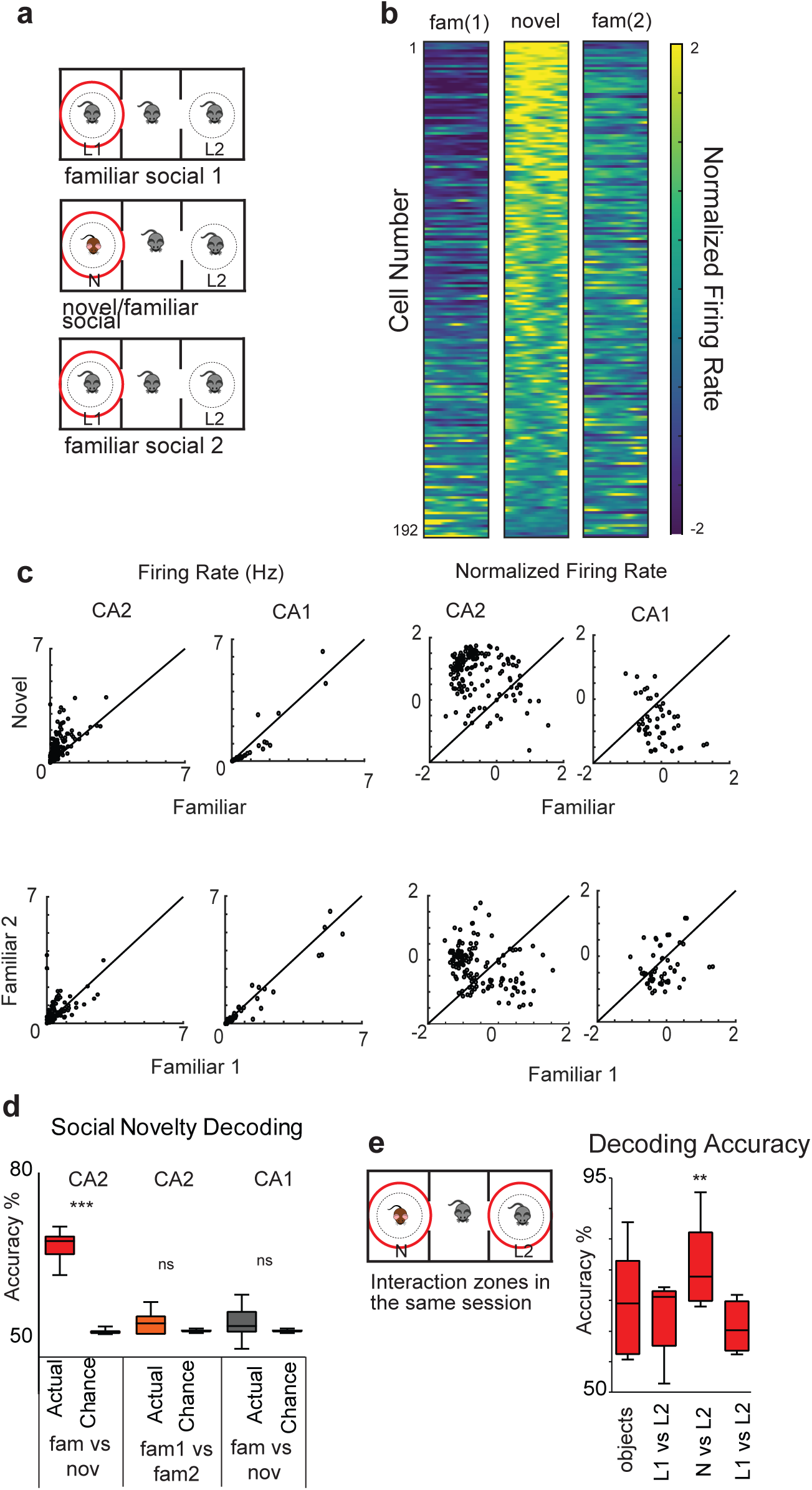
CA2 encoding of social information. (a) CA2 firing rate measured when the animal was within a 7 cm interaction zone around the cup that contained the novel animal. Same physical interaction zone used for all three social sessions (example shows the case in which the novel animal was in the left cup). (b) Mean z-scored firing rates during all social interactions in each of the three social sessions as function of time in interaction zone (192 neurons from 6 mice). Total Interaction time was divided into 50 time bins and firing rate was calculated for each bin to visualize a neuron’s activity. (c) Plot of CA2 and CA1 neuron raw mean firing rates (left graphs) and z-scored mean firing rates (right graphs) in the interaction zone around the novel mouse versus the familiar littermate (the latter was averaged across the two familiar mouse sessions). Each point is a separate cell. (d) An SVM with a linear kernel trained on CA2 activity in the interaction zone performed significantly above chance in decoding interactions with a novel mouse (data from session 4) versus interactions with the familiar mouse (data from sessions 3 and 5; p < 0.0001, Wilcoxon rank-sum test; n=192 neurons from 6 mice). The CA2-based decoder failed to distinguish interactions with the same familiar mouse in session 3 (fam1) versus session 5 (fam2). A decoder based on CA1 activity failed to distinguish interactions between the novel and familiar mouse. (e) Accuracy of a linear decoder trained to distinguish left from right interaction zones in the object, fam1, novel, and fam2 sessions of three-chamber task. Left-right decoding accuracy was above chance for all sessions (p< 0.01, Wilcoxon rank-sum test) and was significantly enhanced when the novel mouse was present (p < 0.01, Wilcoxon rank-sum test, n=192 neurons from 6 mice).

In contrast to the enhanced firing to social novelty, individual CA2 neuron firing rates around the same familiar mouse in session 3 compared to session 5 did not differ significantly (Figure 4b,c). Moreover, only a small fraction of cells showed a z-scored firing rate difference greater than 2 to the same familiar animal (5/192), similar to that predicted by chance for a normal distribution. Furthermore, there was also no significant difference in the two firing rate vectors to the same familiar animal (p>0.05, Wilcoxon rank-sum test). These results indicate that the increased firing to a novel mouse compared to a familiar mouse measured across different sessions was not simply due to the passage of time or CA2 variability (as the difference in time between the two familiar mouse sessions was twice that in the novel versus familiar mouse sessions). Finally, the increase in firing rate around the novel animal was not observed for CA1 neurons under the same conditions (Figure 4c, Supplemental Figure 4), confirming results from Rao and colleagues^30^. Thus, not only does CA2 firing respond to a social stimulus, but CA2 firing is further enhanced in the presence of social novelty. Moreover, this effect is subfield specific, consistent with the importance of dorsal CA2^11, 31^ but not dorsal CA1^13^ in social memory formation.

Is the increase in CA2 firing specific during interactions with a novel social stimulus or does CA2 firing also increase during interactions with a novel object? As the wire cup cages represented novel objects to the mice, we asked whether CA2 firing rates were higher when a mouse was within the interaction zone surrounding the cups in session 2 compared to when the mouse occupied the same location in the empty arena session 1. We did observe a small but significant increase in CA2 z-scored firing rate as mice explored around the novel objects (0.40 ± 0.06) compared to the empty arena (−0.48 ± .05; p<0.01, Wilcoxon rank-sum test). However, the increased firing around the novel animal (0.83 ± 0.06) was significantly greater than that around the novel object (p=0.009, Wilcoxon rank-sum test; Supplemental Figure 4). Thus, although CA2 firing rate increased to both social and non-social novel stimuli, the response was significantly greater for social novelty.

To explore further the social information content in CA2 firing, we asked whether a linear decoder could detect the presence of a familiar mouse versus a novel mouse based on CA2 activity in the same interaction zone around the novel animal in session 4 and the familiar animal in session 3 and 5 that occupied the same cup containing the novel animal (Figure 4d). Indeed, CA2 population activity accurately decoded social interactions with the novel versus familiar mouse among the three social sessions. In contrast, social novelty could not be decoded from CA1 population activity. Importantly, CA2 activity failed to distinguish interactions with the same familiar mouse in session 3 compared to session 5. This further confirmed that CA2 responses to specific social stimuli were responsible for driving decoder performance as opposed to a purely time-dependent change in CA2 firing patterns over the various sessions.

As an additional probe of CA2 information content, we determined whether a decoder could discriminate in a single session whether a subject mouse was exploring within the interaction zone around the cup in the left chamber versus the cup in the right chamber, a comparison that can incorporate both spatial and non-spatial cues (Figure 4e). In all four sessions that contained the identical cups (sessions 2-5), the decoder was able to determine whether an animal was exploring within the left versus right interaction zones at a level significantly better than chance. The ability of CA2 firing to decode left from right with two identical objects present suggests that CA2 firing may contain coarse spatial information (although it is possible that decoder performance is driven by subtle physical differences in the nominally “identical” objects). Of interest, left-right decoder performance was significantly enhanced in the novel-familiar mice session compared to either the object session or the two familiar mice sessions (Figure 4e), supporting the view that CA2 firing contained significant information on social novelty.

### CA2 neurons in *Df(16)A^+/-^* mice showed altered spatial, contextual and social firing

If the firing properties of CA2 neurons are relevant for social cognition, we might expect CA2 activity to be altered in models of neuropsychiatric disease with known deficits in social memory. To test this possibility, we recorded the activity of 128 CA2 neurons during the three-chamber task from five *Df(16)A^+/-^* mice (Figure 5a; Supplemental Table 1, Supplemental Figure 5). Such mice were previously found to have a deficit in social memory that is associated with a late developmental decrease in density of PV^+^ inhibitory neurons within CA2, resulting in decreased CA2 synaptic inhibition^22^. CA2 pyramidal neurons of these mice also displayed a negative shift in their resting potential as a result of an increase in TREK-1 K^+^ channel resting current^22^. We therefore also examined whether any alterations in CA2 firing and/or social memory deficit in these mice could be rescued by the selective TREK-1 peptide antagonist spadin^32^, using a different group of *Df(16)A^+/-^* mice (91 cells from 5 mice).

**Figure 5.**
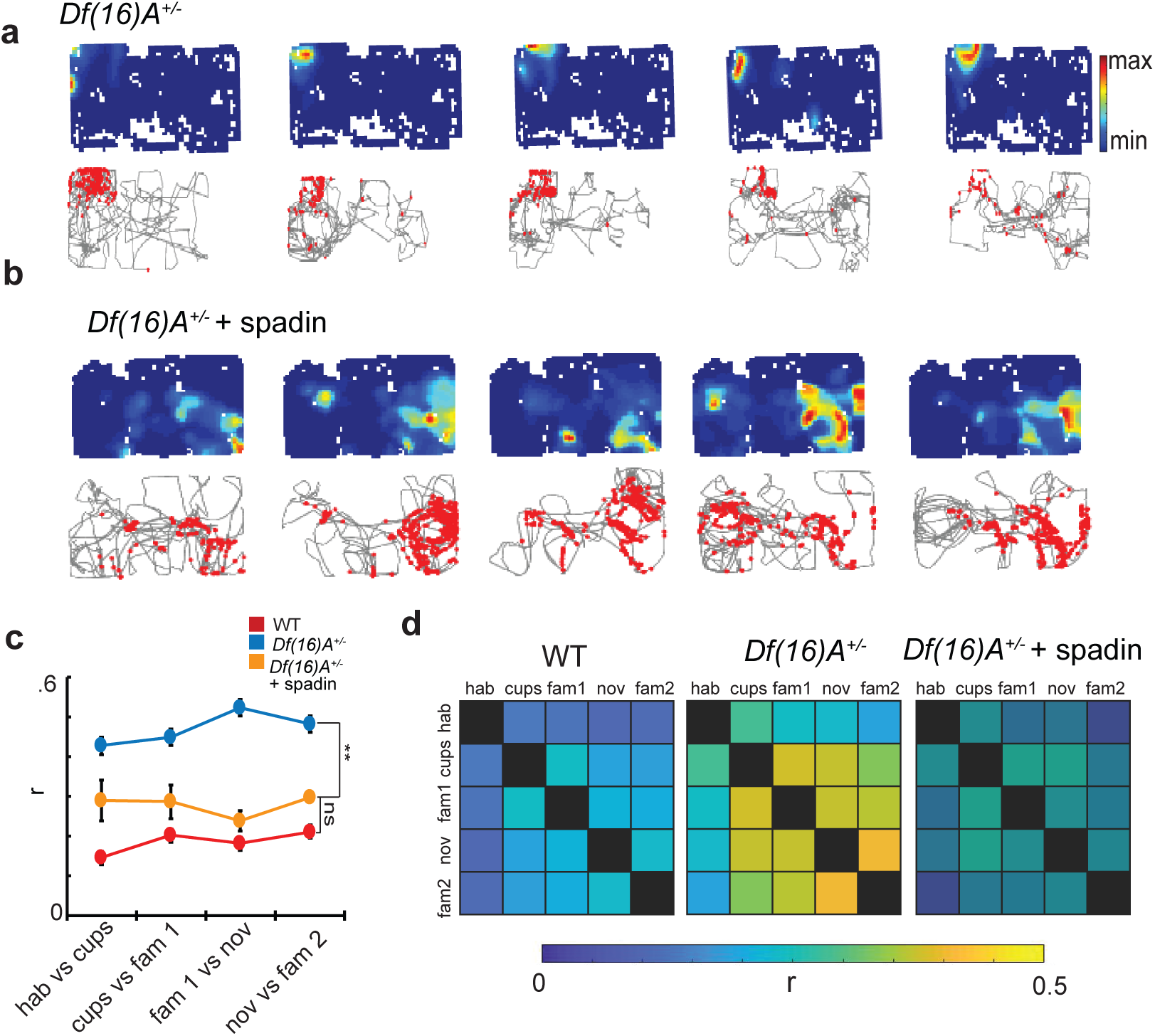
Spatial firing of CA2 neurons in Df(16)A+/- mice and effect of systemic injection of TREK-1 antagonist spadin. (a) Example CA2 spatial firing across the 5 sessions of the three-chamber task from Df(16)A+/- mice. CA2 example neuron spatial firing (maximum firing rates in sessions 1-5 were: 5, 2, 4, 2, and 2 Hz). (b) CA2 example neuron firing from a Df(16)A+/- mouse 30 min after injection of spadin (maximum firing rates in sessions 1-5 were: 2, 7, 17, 2, and 12 Hz). (c,d) Place field stability between pairs of consecutive sessions (c) and between all sessions (d) in wild-type mice (same data as in Figure 2) compared to Df(16)A+/- mice in the absence (n=128 neurons from 5 mice) and presence (n=91 neurons from 5 mice) of spadin.

We found that the mean firing rate of CA2 neurons from *Df(16)A^+/-^* mice during the five sessions of the three-chamber task was significantly decreased compared to that in two groups of wild-type mice, unrelated wild-type mice of the same C57Bl/6J genetic background and wild-type littermates (Supplemental Figure 6b). This suggests that the inhibitory effect of CA2 pyramidal neuron hyperpolarization may predominate over the excitatory effect of decreased feedforward inhibition. Surprisingly, we also found that the spatial coding properties of CA2 neurons in the mutant strain were significantly enhanced so that they more closely resembled the spatial coding characteristic of CA1 pyramidal cells (Figure 5, Supplemental Figure 5). Thus, CA2 neuron spatial firing in *Df(16)A^+/-^* mice showed a significant increase in stability across the sessions of the three-chamber task in (Figure 5c,d). Moreover, the CA2 PNs had fewer and larger place fields with a higher average spatial information and selectivity compared to wild-type mice (Supplemental Figure 5c-f). The increase in place field stability was not due to the increase in field size, as the y-axis intercept of the regression line for a plot of field size versus stability^33^ was significantly higher for *Df(16)A^+/-^* animals (*Df(16)A^+/-^* = 0.29 ± 0.032; wild-type = 0.15 ± 0.028; p < 0.001, ANCOVA).

As a further indication of improved spatial coding, we found that spatial location could be determined at a level significantly better than chance by a linear decoder trained on CA2 PN population activity in the *Df(16)A^+/-^* mice (Figure 6a,b), in contrast to the poor decoder performance based on CA2 activity in wild-type mice (Figure 2d). Finally, although spatial decoding abilities of CA2 firing in *Df(16)A^+/-^* mice was enhanced, the ability of CA2 activity to decode session was significantly impaired (Figure 6c,d), implying a deficit in contextual coding.

**Figure 6.**
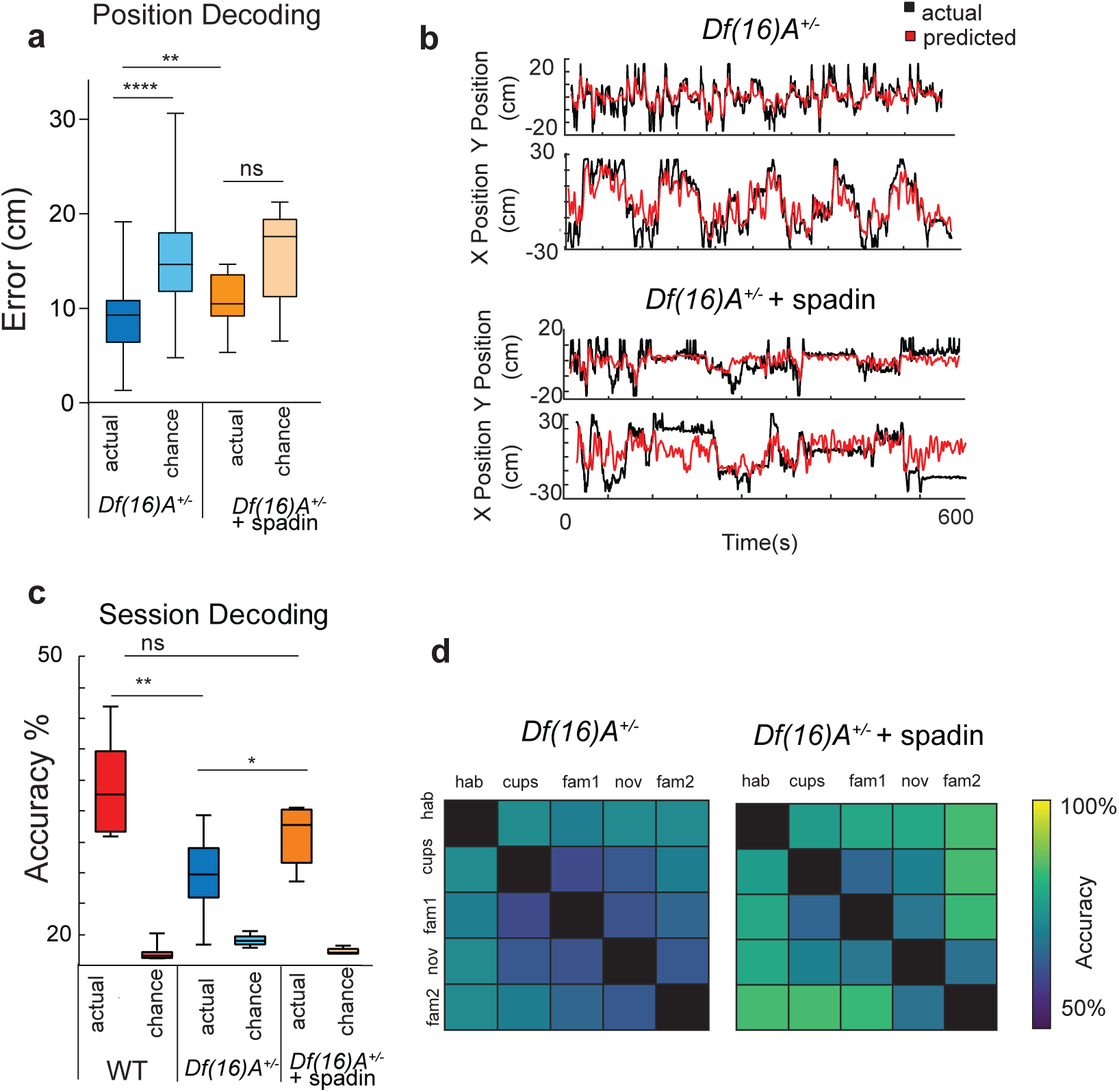
CA2 population activity decoding of position and session in Df(16)A+/- mice and the effect of spadin. . (a,b) Position of the animal was decoded significantly better than chance (p < 0.001, Wilcoxon rank-sum test) from CA2 population activity in Df(16)A+/- mice (n=128 neurons from 5 mice). Spadin decreased decoding accuracy (p < 0.01, Wilcoxon rank-sum test) to chance levels (n=91 neurons from 5 mice). (c) Overall CA2 decoding accuracy for session in three-chamber task (context) is impaired in Df(16)A+/- mice compared to wild-type mice (p < 0.01, Wilcoxon rank-sum test), although it is significantly greater than chance (p < 0.05, Wilcoxon rank-sum test). Treatment with spadin significantly increased session decoding performance (p < 0.01, Wilcoxon rank-sum test). (d) Decoding accuracy for pairs of sessions in absence and presence of spadin.

Are the social coding properties of CA2 neurons also altered in the *Df(16)A^+/-^* mice? Indeed, we found a significant impairment in the ability of CA2 activity from these mice to encode social information and social novelty (Figure 7). Thus, CA2 neurons of *Df(16)A^+/-^* mice failed to show an increase in firing around a novel social stimulus (Figure 7b,c; compare with Figure 4b,c). In addition, the CA2 population normalized firing rate around the novel mouse in session 4 was did not differ from the population firing rate around the familiar mouse in either session 3 or 5 (Figure 7b,c; p>0.05, Kruskal Wallis). This is in distinction to the significant difference in firing rate vectors we observed for wild-type mice when exploring a familiar versus novel mouse (Figure 4b,c). In addition, only 3/128 CA2 PNs showed a significant increase (>2 SD) in normalized firing rate when interacting with a novel animal compared to a familiar one (Figure 7c), in contrast to the 20% of cells in wild-type mice that increased their firing rate in response to social novelty (Figure 4b,c).

**Figure 7.**
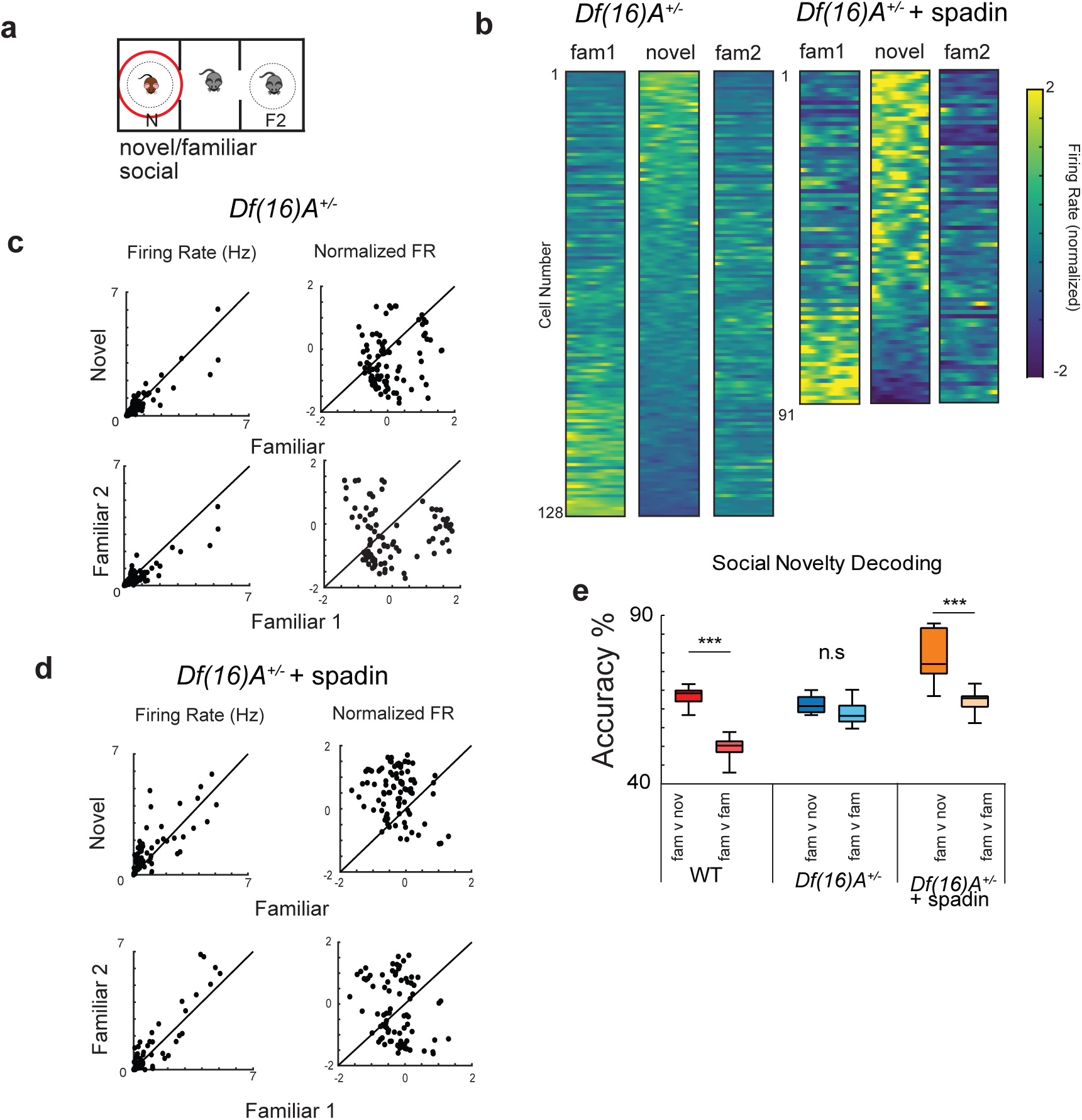
Social coding deficits in Df(16)A+/- mice and its escue by spadin. (a) CA2 firing was analyzed in the interaction zone defined by the cup containing the novel animal in three social sessions. (b) Z-scored CA2 firing rates for each cell during social sessions as a function of time in the interaction zone in untreated (left, n=128 neurons from 5 mice) and spadin-treated (right, n=91 neurons from 5 mice) Df(16)A+/- mice. (c),(d) Comparison of CA2 neuron firing rates in Df(16)A+/- animals in interaction zone around the novel versus familiar mouse (top) or during interactions with the same familiar mouse in session 3 versus 5 (bottom). Graphs on left show mean firing rates (Hz); graphs on right show mean z-scored firing rates. (c) Firing in absence of drug; (d) Firing in presence of spadin. (e) Accuracy by which CA2 activity decoded interactions with the familiar versus novel mouse (fam v nov) and with same familiar mouse in sessions 3 versus 5 (fam v fam). Data shown for wild-type mice (same as Figure 3d) compared to Df(16)A+/- mice in absence and presence of spadin. Decoding for novel versus familiar social stimuli was significantly greater than decoding for the same familiar mouse in wild-type and spadin-treated Df(16)A+/- mice (p < 0.001, Wilcoxon rank-sum test), but not in untreated Df(16)A+/- mice (p > 0.05, Wilcoxon rank-sum test).

Next we examined social coding properties of CA2 in *Df(16)A^+/-^* mice by training a linear decoder on CA2 population activity during periods of exploration within the same interaction zone around a novel or a familiar mouse, as described above. Although the decoder was able to distinguish interactions with the novel versus familiar mouse, decoder performance was only barely above chance levels (p = 0.04, Wilcoxon rank-sum test). Importantly, the decoder showed no improvement in its ability to discriminate between interactions with the novel versus familiar mouse (session 4 versus sessions 3 and 5) compared to its ability to discriminate between interactions with same familiar mouse in sessions 3 versus 5 (Figure 7e; p=0.36, Wilcoxon rank-sum test). This contrasts with the effect of social novelty to significantly enhance decoder performance in wild-type mice (Figure 4d). Thus, CA2 activity in the *Df(16)A^+/-^* mice had a reduced response to a social stimulus and social novelty, consistent with the deficit in social memory of these mice.

### TREK-1 inhibition rescued social memory and CA2 social coding deficits in *Df(16)A^+/-^* mice

To what extent can the changes in CA2 firing in the *Df(16)A^+/-^* mice be rescued by inhibition of TREK-1 K^+^ current^21^? Do the alterations in CA2 firing observed above actually contribute to the social memory deficit of the *Df(16)A^+/-^* mice? We addressed these questions by investigating the effects on CA2 firing properties and social memory behavior of systemic injection of spadin, a naturally occurring selective peptide antagonist of TREK-1^31^. We found that the changes in CA2 spatial, contextual and social firing properties were largely rescued when *Df(16)A^+/-^* mice were injected intraperitoneally with 0.1 ml of 10^-5^ M spadin 30 min prior to testing (Figures 5-7; Supplemental Figure 5b-f). In contrast, injection of a control group of *Df(16)A^+/-^* mice with saline, the spadin vehicle, had no effect on CA2 firing (Supplemental Figure 6).

Spadin administration increased CA2 neuron mean firing rate throughout the five sessions of the three-chamber task to wild-type values (Supplemental Table 1, Supplemental Figure 5b), consistent with the idea that the decreased firing rate in *Df(16)A^+/-^* mice was caused by TREK-1 upregulation. Compared to untreated *Df(16)A^+/-^* mice, CA2 neurons in spadin-treated *Df(16)A^+/-^* mice had more plentiful but smaller place fields (Figure 5b; Supplemental Figure 5c,d) that were less stable across sessions (Figure 5c,d), properties more similar to CA2 firing properties in wild-type animals. Notably, spadin also decreased the spatial selectivity and information content of CA2 PN firing (Supplemental Figure 5e,f, Supplemental Table 1). In addition, spadin decreased position decoding performance in the *Df(16)A^+/-^* mice to chance levels (Figure 6a,b), as found for wild-type mice (Figure 2d). In contrast to its effects to degrade CA2 spatial information coding, spadin enhanced the ability of CA2 population activity to decode three-chamber task session in which a mouse was engaged, thus rescuing CA2 contextual coding (Figure 6c,d).

Does TREK-1 antagonism with spadin also rescue the social coding properties of CA2? Indeed, following spadin treatment CA2 firing in *Df(16)A^+/-^* mice around a novel mouse was now significantly greater than firing around a familiar mouse (Figure 7a-d), with 33/91 cells showing a significant (>2 SD) increase (Figure 7d), similar to the findings in wild-type mice (Figure 4). Moreover, in the presence of spadin, the CA2 population firing rate vector around the novel versus familiar mice differed significantly (p,0.01, Wilcoxon rank-sum test), similar to our findings in wild-type mice but in contrast to uninjected or saline-injected *Df(16)A^+/-^* mice (Figure 7b,d; Supplemental Figure 6). Finally, spadin injection rescued the normal finding that decoder performance in discriminating a novel from a familiar mouse was greater than decoder performance in discriminating the same familiar mouse in session 3 versus session 5 (Figure 7e).

### Systemic and CA2-selective TREK-1 inhibition rescues social memory in *Df(16)A^+/-^* mice

If the social memory deficit in the *Df(16)A^+/-^* mice was related to impaired CA2 social coding, we would expect that spadin should also rescue social memory given its ability to rescue CA2 social coding. We first explored social memory performance using a direct interaction test (Figure 8a-d). In this test, a subject mouse was first exposed to a novel stimulus mouse for 2 min in trial 1. The mice were then separated for 30 min and the subject mouse was then re-introduced to the now familiar stimulus mouse for 2 min in trial 2. In wild-type mice social memory is normally expressed as a decrease in interaction time with the stimulus mouse in trial 2 relative to trial 1, reflecting the decrease in social novelty. Whereas saline-treated *Df(16)A^+/-^* mice showed no decrease in social exploration in trial 2, indicative of a loss of social memory, *Df(16)A^+/-^* mice treated with spadin showed a significant decrease in social interaction time in trial 2, similar to that seen in wild-type mice treated with saline or spadin (Figure 8a-d). Importantly, spadin-treated *Df(16)A^+/-^* mice showed no decrease in interaction time when a second novel mouse was introduced in trial 2, showing that the decrease in interaction to the same mouse encountered in trial 1 was due to decreased social novelty associated with social memory, and not simply task fatigue (Supplemental Figure 7b). We also found that spadin rescued social memory performance during *Df(16)A^+/-^* mouse social interactions in the three-chamber task (Supplemental Figure 7d-f).

**Figure 8.**
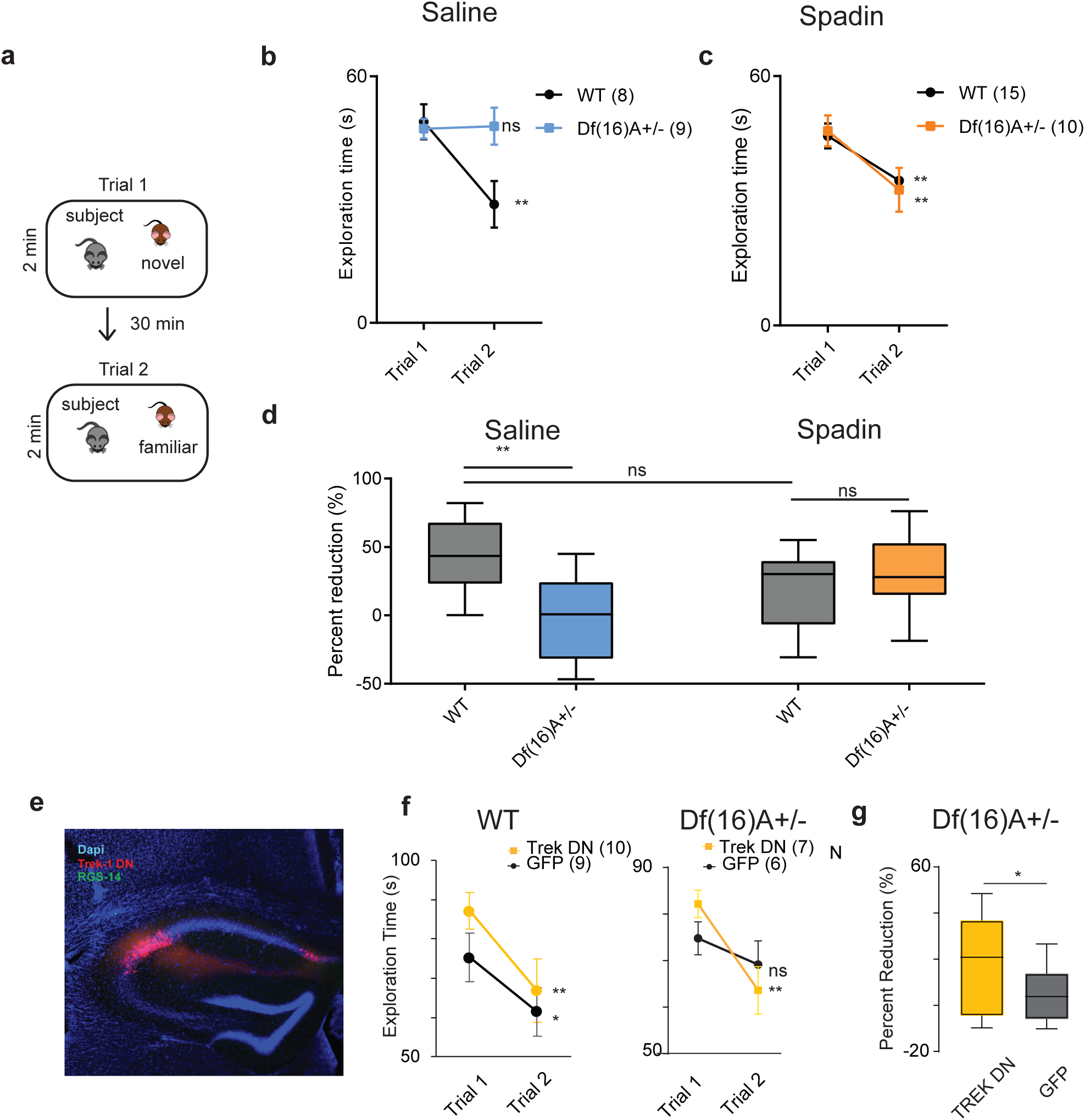
Effect of TREK-1 inhibition onsocial memory deficits in Df(16)A+/- mice. (a) The direct interaction task. Trial 1, A subject mouse was presented with a novel stimulus mouse for 2 min. The novel mouse was then removed from the cage. Trial 2, After 30 min the same (now familiar) stimulus mouse is reintroduced. (b) Wild-type mice injected with saline showed a decreased interaction time in trial 2 compared to trial 1 (p< 0.01, paired t-test, n=8 mice). Df(16)A+/- mice injected with saline showed no decrease in interaction time (p>0.05, n=9 mice). (c) In mice injected with spadin 30 min prior to trial 1, interaction time in trial 2 compared to trial 1 decreased significantly for both wild-type (p< 0.01, paired t-test, n=15 mice) and Df(16)A+/- mice (p < 0.01, paired t-test, n=10 mice). (d) Percent reduction in interaction time is significantly lower in saline-treated Df(16)A+/- mice than other experimental groups (ANOVA, p=0.009). Spadin-treated Df(16)A+/- mice do not differ from saline- or spadin-treated wild-type mice (p=0.37, paired t-test). (e) Immunohistochemistry showing viral mediated expression in CA2 (identified by CA2 marker RGS-14, green signal) of TREK-1 DN tagged with GFP (TREK-1 DN, red signal). (f,g) Social memory in wild-type and Df(16)A+/- mice expressing TREK-1 DN or GFP (control) in CA2. There was a significant decrease in interaction time in trial 2 for wild-type mice expressing TREK-1 DN (p=0.005, n=10 mice) or GFP (p=0.03, n=9 mice) and for Df(16)A+/- mice expressing TREK-1 DN (p=0.001, n=7 mice) but not GFP (p=0.07, n=6 mice; paired t-tests for all comparisons).

Because the effects of spadin were observed following systemic injection, it was important to determine whether its behavioral effect resulted from a specific action in CA2 or was mediated by effects on some other brain region. We therefore we injected a TREK-1 dominant negative virus (Trek-1 DN)^34^ in CA2 of *Df(16)A^+/-^* and wild-type mice to decrease TREK-1 K^+^ current selectively in this region. To further limit expression to CA2 we used AAV2/5, whose serotype causes it to have a natural tropism to infect CA2 compared to neighboring CA1 or CA3 regions^35^. We found that *Df(16)A^+/-^* animals expressing TREK-1 DN in CA2 showed a significant improvement in social memory in the direct interaction test, manifest as a decreased social exploration of the now-familiar stimulus mouse in trial 2, as compared to control *Df(16)A^+/-^* animals expressing GFP (Figure 8g-i). Moreover, mice expressing TREK-1 DN showed no decrease in interaction time between trial 1 and trial 2 when a novel animal was introduced on trial 2, indicating that the decreased interaction time to the familiar mouse in trial 2 was not due to fatigue (Supplemental Figure 7c). As a further control, we found that injection of the TREK-1 DN virus in CA2 did not alter social memory performance in wild-type mice (Figure 8f).

## Discussion

Here we report that the firing of dorsal CA2 pyramidal neurons, which play a critical role in social memory, was enhanced during an animal’s interactions with a novel compared to a familiar conspecific. At the population level, CA2 activity successfully decoded both social and non-social contexts and discriminated between interactions with a novel versus a familiar mouse. These changes in firing could reflect coding of social novelty or the increased salience of a novel social stimulus compared to a familiar social stimulus. As novel conspecifics are generally extremely salient stimuli, deficits in encoding of salience could also lead to social behavioral deficits. While we found evidence for highly significant social and contextual coding in CA2, the same neurons provided at best a weak representation of spatial information at either the single cell or population level, especially compared to their neighboring CA1 neurons. Although most discussions of hippocampal firing focus on location-based measures, our results indicate that this fails to capture the most salient aspects of CA2 firing, especially when contextual elements in an environment are changed, with the presence of a novel conspecific eliciting the strongest response.

Although CA2 activity was clearly responsive to social stimuli, our experiments were not designed to reveal whether the firing of individual CA2 neurons or the CA2 population contained a representation for a social engram that encodes the social identity of a familiar conspecific, enabling an animal to distinguish one familiar conspecific from another. Okuyama et al (2016) reported that a subclass of neurons in ventral CA1 that project to the shell of the nucleus accumbens do form a social engram, with about 10% of neurons selectively active during interactions with a specific familiar mouse. Moreover, these authors found that such neurons were necessary and sufficient to encode and retrieve a social memory. Unlike our results in dorsal CA2, ventral CA1 neurons were not reported to increase their firing in response to social novelty. This is perhaps surprising as our laboratory found that dorsal CA2 provides excitatory input to the same subset of ventral CA1 that Okuyama et al identified and that this CA2 input is necessary for encoding social memory^31^. How the novel social coding in dorsal CA2 is transformed into familiar specific firing in ventral CA1 remains an open question, although the dorsal CA2 inputs do recruit substantial feedforward inhibition in ventral CA1.

Evidence in support of an important behavioral role of the CA2 social firing properties came from our analysis of the *Df(16)A^+/-^* mouse model of the human 22q11.2 deletion syndrome. Consistent with the profound deficit these mice exhibit in both contextual fear memory^36^ and in social memory^22^, we found that the firing of CA2 pyramidal neurons in these mice show diminished contextual and social responses and have a reduced ability to decode context or social novelty. Surprisingly, we found that CA2 activity in these mice had improved spatial encoding properties, including increased place field stability and spatial selectivity that was associated with an improved ability to decode spatial location.

At the cellular level, CA2 neurons in the *Df(16)A^+/-^* mice were previously found to have decreased feedforward synaptic inhibition in response to activation of their CA3 Schaffer collateral inputs and a more negative resting potential compared to control mice^22^. The latter effect was proposed to result from an upregulation of the resting K^+^ current carried by TREK-1 channels^22^, whose mRNA expression is normally highly enriched in CA2 relative to other hippocampal regions^37^. These two cellular effects should have opposing actions on CA2 firing, with decreased inhibition enhancing and hyperpolarization suppressing CA2 activity. Our finding that mean CA2 firing rate was significantly decreased in the mutant mice indicates that the net effect of the cellular alterations may be dominated by membrane hyperpolarization. Consistent with this view, we found that decreasing TREK-1 current by injection of the TREK-1 antagonist spadin or injection of a TREK-1 DN virus in the *Df(16)A^+/-^* animals resulted in a rescue of their social memory deficits.

At present, the molecular mechanisms linking the loss of genes found in the microdeletion to the upregulation of TREK-1 current remain unknown. It is possible that certain gene products in the locus could normally suppress TREK-1 expression, for example as found for the effect of micro RNA 185, which leads to the derepression of its gene target *Mirta22*^38^. Alternatively, as TREK-1 channel activity is regulated by a number of intracellular signaling cascades^39^ the effects on TREK-1 current could be indirect as a result of altered CA2 firing due to decreased inhibition or altered activity in a modulatory input to CA2.

Why should selective TREK-1 inhibition produce an effective rescue of both social memory behavior and CA2 firing properties in the *Df(16)A+/-* mice given that TREK-1 antagonism is not expected to restore the normal level of synaptic inhibition? The two major excitatory inputs to dorsal CA2 pyramidal neurons come from entorhinal cortex layer II stellate cells through the perforant path and from hippocampal CA3 pyramidal neurons via the Schaffer collaterals, although the net excitatory action of the latter inputs is normally suppressed by strong feedforward inhibition. Of interest, Piskorowski and colleagues^22^ found that the decrease in CA2 feedforward inhibition in the *Df(16)A^+/-^* mice was selective for the Schaffer collateral inputs, with no change in feedforward inhibition through the entorhinal cortical inputs. Thus, if CA2 were to normally receive its major social information from the entorhinal cortical inputs as opposed to CA3, the rescue of CA2 neuron hyperpolarization could well be sufficient to restore normal levels of CA2 social information processing. That social information may arrive via the direct cortical inputs as opposed to CA3 is consistent with a recent study showing that silencing dorsal CA3 did not affect social memory^40^.

To our knowledge, our results provide the first instance of a mechanism-based pharmacological rescue of the social deficits in a mouse genetic model of schizophrenia. This finding is notable given the difficulty in treating the negative symptoms of schizophrenia, including social withdrawal. Spadin administration also rescued the social coding deficits seen in *Df(16)A^+/-^* CA2 activity. Interestingly, spadin administration, in addition to rescuing social and contextual coding, reverted the spatially selective firing properties of CA2 neurons to the less precise spatial firing characteristic of CA2 in wild-type mice. This suggests that the increased spatial stability in the *Df(16)A^+/-^* mice may actual contribute to impaired social coding by altering the normal mixed selectivity of CA2 encoding to social, spatial, contextual and temporal signals to a more selective coding mode dominated by spatial information.

The apparent gain-of-function of improved CA2 spatial coding in the *Df(16)A^+/-^* mice was surprising, although pathological hyperstability of place fields has also been described in the *Fmr1* KO mouse model of fragile X syndrome^41^. Moreover, the *Df(16)A^+/-^* mice were previously found to have deficits in reward-related remapping of CA1 place fields^42^. The more stable spatial firing in CA2 of the *Df(16)A^+/-^* mice may reflect a deficit in remapping in response to altered context. Perhaps the pyramidal cell hyperpolarization may render CA2 neurons less sensitive to weak excitatory or neuromodulatory inputs that convey contextual information. In addition, the loss of CA2 feedforward inhibition through the Schaffer collateral input may shift the balance of excitatory input to favor the more spatially oriented information conveyed by CA3. Finally, as CA2 is enriched in receptors for the social neuropeptides oxytocin^43^ and vasopressin^44–46^, improper integration of these social signals could contribute to the behavioral and social coding deficits seen in the *Df(16)A^+/-^* mice.

Our results provide further support that CA2 and its malfunction contributes importantly to normal social behavior and to social behavioral abnormalities characteristic of certain neuropsychiatric disorders, including schizophrenia. Moreover, our findings emphasize the potential importance of CA2 and TREK-1 as a target for novel therapeutic approaches to treating social endophenotypes associated with these disorders. Finally, the strong and consistent correlation we observe between CA2 firing properties and social memory behavior in wild-type mice, *Df(16)A^+/-^* mice, and *Df(16)A^+/-^* mice treated with spadin provides additional support for the view that CA2 social firing properties contribute to the role of this region in the encoding, storage and recall of social memory.

## Acknowledgements

We would like to thank Joseph Gogos for initially providing the Df(16)A^+/-^ mice and for ongoing advice and guidance. We thank Yuichiro Matsushita of Ono Pharmaceuticals for suggesting we study the action of spadin. We also thank Y. Mara Zafrina, Ambar Kleinbort, and Hannah G. Yueh for their technical support, Bina Santoro for assistance with and animal breeding, and C. Daniel Salzman and Dmitry Aranov for helpful discussions and comments on the manuscript. This work was supported by a grant from NSF GRFP to M.L.D., grants R01MH104602 and R01MH106629, S.A.S, P.I., support from the Zegar Family Foundation, and a grant from Ono Pharmaceuticals.

## Author Contributions

M.L.D, J.A.G, S.F, and S.A.S designed the experiments and analyses. M.L.D. performed the in vivo recordings and M.L.M and T.M. did the behavioral experiments. M.L.D and F.S. analyzed the data. M.L.D and S.A.S wrote the manuscript.

## Data and Code Availability Statement

The datasets generated during and/or analysed during the current study are available from the corresponding author on reasonable request. All scripts for analyzing data are also available by request.

## Methods

We bred *Df(16)A^+/-^* mice and their wild-type littermates on a pure (>99.9%) C57BL/6J background (The Jackson Laboratory) as previously described^23^. Experiments were carried out on adult male mice (22-28 g, 3-6 months old). Mice were housed 3-5 in a cage under a 12:12 h light/dark cycle with access to food and water *ad libitum*. Experiments were conducted during the light cycle. All procedures were approved by the Animal Care and Use Committee of Columbia University and were in accordance with the National Institutes of Health guidelines for care and use of animals.

### Surgical Procedures

19 mice (10 *Df(16)A^+/-^* mice, 3 wild-type littermates, 6 wild-type C57Bl/6J non-littermates), were implanted with electrode bundles containing 7-8 tetrodes in a moveable drive using sterile surgical techniques. Wild-type littermates and wild-type non-littermates were not significantly different in any of the physiological measures discussed with the exception of spatial stability, in which littermates were significantly less stable that non-littermate wild-type mice; both control groups were significantly less stable than both CA1 and *Df(16)A+/-* CA2 recordings (Supplemental Figure 8). Animals were anesthetized with 2-5% isoflourane and placed in a stereotaxic frame. Craniotomies were made above CA2 (1.8 mm posterior to bregma, 2.15 mm lateral to the midline, ∼1.5 mm below the brain surface) or CA1 (1.9 mm posterior to bregma, 1.8 mm lateral to the midline, ∼1 mm below the brain surface). To prevent damage to the recording site, tetrodes were implanted above the structure and turned down 50-150 μm per day after recovery from surgery. A skull screw was placed over the contralateral visual cortex to serve as ground. To verify recording locations, 50 μA of current was passed through the channels at the end of the experiment to create an electrolytic lesion (Supplemental Figure 9). For viral injections, 200 μl of either AAV2.5-hsyn-mCherry or AAV2.5.-hsyn-TREK-1DN-GFP were injected bilaterally into CA2 (1.8 mm posterior to bregma, 1.8 mm lateral to the midline, 1.2 mm below the brain surface). After experiments animals were perfused, and brains were cut on a vibrotome and stained for RGS-14 and NeuN. One animal was excluded from the TREK-1 DN group because there was only unilateral expression of the virus..

### Recording and Spike Sorting

Recordings were amplified, band-pass filtered (1–1,000 Hz LFPs, 300–6,000 Hz spikes), and digitized using the Neuralynx Digital Lynx system or the OpenEphys GUI. LFPs were collected at a rate of 2 kHz, while spikes were detected by online thresholding and collected at 30 kHz. Units were initially clustered using Klustakwik, sorted according to the first two principal components, voltage peak and energy from each channel. Clusters were then accepted, merged or eliminated based on visual inspection of feature segregation, waveform distinctiveness and uniformity, stability across recording session, and inter-spike interval distribution.

### Behavior: Three-chamber Interaction Task

Mice were given 1 week to recover from surgery, after which tetrodes were turned down to stratum pyramidale of the hippocampus. After tetrodes reached the hippocampus and were stable for at least 48 hours (Supplemental Figure 9), animals were habituated to the three-chamber arena (60 × 40 cm) for 40-50 minutes. Barriers were placed in the environment every 10 minutes to briefly isolate mice in the center chamber to match the protocol of the three-chamber task (Figure 1d). The following day animals were run in the three-chamber social interaction task shown in Figure 1a. The subject mouse was isolated in the central chamber between each of the five sessions by placement of barriers. Side chambers were quickly wiped with 70% alcohol between each session to rid the side chambers of any olfactory cues from the previous session. The trajectory of the animal was recorded in Neuralynx using LEDs on the head to track the position of the head or using custom MATLAB software for tracking webcam images. Trajectory and behavior were analyzed using custom scripts in MATLAB. Interaction zones were defined as a 7 cm annulus from the edges of each respective cup. Experimenters were blind to experimental condition during recordings.

### Behavior: Direct Interaction Task

The direct interaction task was performed on 9 wild-type mice and 9 *Df(16)A^+/-^* mice injected with saline control and with 16 wild-type and 13 *Df(16)A^+/-^* mice injected with spadin. Subject mice were habituated to the cage for 30 min. A novel juvenile male mouse was placed into the cage for 2 min (Trial 1), during which the subject mouse was allowed to explore the juvenile. Mice that interacted for less than 24 seconds in Trial 1 were excluded from analysis (1 wild-type mouse was excluded from the saline group; 1 wild-type and 3 *Df(16)A+/-* mice were excluded from the spadin group). The juvenile mouse was removed for 30 minutes and then placed back into the cage with the subject for an additional 2 minutes (Trial 2). For the novel-novel version of the direct interaction task (Supplemental Figure 7), a second novel juvenile mouse was placed in the cage in Trial 2. Behavioral videos were recorded and analyzed in Anymaze 2.0; social interactions were defined as periods of facial or anogenital sniffing, grooming of the juvenile mouse, and periods of chasing the juvenile. Researchers were blind to experimental condition during both the behavioral experiments and the analysis.

### Spatial Fields

Spatial analyses were performed with custom written scripts in MATLAB. From each session, X,Y positions from LEDs placed on the animals head during the three-chamber were projected onto the apparatus axis. The position and spiking data were binned into 5-cm wide segments, generating the raw maps of spike number and occupancy probability, with unvisited bins for each session represented as NaNs. Rate map, number of place fields, field sizes, spatial information, and selectivity were calculated. A Gaussian kernel (SD = 5.5 cm) was applied to both raw maps of spike and occupancy, and a smoothed rate map was constructed by dividing the smoothed spike map by the smoothed occupancy map. A place field was defined as a continuous region, of at least 9 cm, where the firing rate was above 10% of the peak rate in the maze, with a peak firing rate >2 Hz. Spatial stability was calculated as the Pearson’s correlation (r) of the firing rate in each binned location for each session only using those spatial bins that were visited in all sessions being compared.

### Statistics and Normalization

All effects presented as statistically significant exceeded an α-threshold of 0.05. All independence tests were two-tailed. All independence testing of paired values (that is, changes across conditions) used paired t-tests or (where stated) signed rank tests. All t-tests and rank tests performed with more than two groups were done post-hoc following ANOVA tests or Kruskal Wallis tests, except where Bonferroni correction for multiple comparisons is specifically cited. Normalization refers to z-scored data. For all box plots displayed the center line is the mean; box limits are upper and lower quartiles; whiskers and min to max values in data sets.

### Position Decoder

For decoding position^47^, we considered the different sessions of the tasks separately to evaluate the different valences. For all the datasets, unless otherwise specified, we used 10-fold cross validation to validate the performance of the decoders. We divided each individual 10 min trial into 10 temporally contiguous periods of equal size in terms of number of data points (spikes). We then trained the decoders using the data from 9 of the 10 periods and tested the performance of the decoder on the remaining data in the session.

To decode the position of the animal, we first divided the arena into 12 × 8 equally sized, square bins. We then labeled each time point with the discrete location in which the animal was found. For each pair of locations, we trained a Support Vector Machine (SVM) classifier with a linear kernel to classify the cell activities into either of the two assigned locations using all the identified cells unless specified otherwise. We used only the data corresponding to the two assigned locations; to correct for unbalanced data due to inhomogeneous exploration of the arena we balanced the classes with weights inversely proportional to the class frequencies^48^. The output of the classifiers was then combined to identify the location with the largest number of votes as the most likely location. The decoding error reported corresponds to the median physical distance between the center of this location and the actual position of the mouse in each time bin of the test set, unless otherwise specified. For datasets with different numbers of cells we randomly down sampled until all groups had equal numbers of cells.

To assess the statistical significance of the decoder, we computed chance distributions of decoding error using shuffled distributions of spike events. Briefly, for each shuffling, we assigned a random time bin to each spike event for each cell independently while maintaining the overall density of spike events across all cells. That is, we chose only time bins in which there were spike events in the original data and kept the same number and magnitude of the events in each time bin. This method destroys spatial information as well as temporal correlations but keeps the overall activity across cells. We trained one decoder on each shuffled distribution and pooled all the errors obtained. We finally assessed the statistical significance of the decoding error for the 5-fold cross-validation of the original data by comparing them to the distribution of errors obtained from the shuffled data using the non-parametric Mann-Whitney U test, from which we obtained a p-value of significance.

### Session/Social Information Decoder

All sessions were divided into 5 equal time bins, and a Support Vector Machine (SVM) was trained to decode either which session the animal was engaged or with which animal the subject mouse was interacting with, using 4 time bins to train and the remaining time bin to test. In the case of unequal interaction times, training sets were subsampled to the length of the shorter interactions. Chance was determined by training and testing the decoder on shuffled data. Statistical significance was assessed by comparing to the distribution of the shuffled data using the Mann-Whitney U test.

### Spadin administration

For the three-chamber experiments described, 0.1 ml of 10^-5^ M spadin (Tocris) or saline was administered intraperitoneally 30 minutes before the three-chamber interaction task in 5 of the *Df(16)A^+/-^* mice. CA2 firing properties in *Df(16)A^+/-^* mice treated with saline did not show any significant differences from the untreated *Df(16)A^+/-^* mice (Supplemental Figure 7). For the direct interaction task described, either 0.1 ml of saline or 0.1 ml of 10^-5^ M spadin was injected into *Df(16)A+/-* mice and wild-type littermate controls 30 minutes before trial 1 of the direct interaction.

### TREK-1 Dominant Negative Virus Generation

From^34^: A dnTREK-1 mutant was created from the mTREK-1 plasmid by the introduction of two point mutations in the selectivity filter of the pore region (G161E and G268E). The mutations were introduced using the QuikChange kit (Stratagene). The primers designed to generate the mutation of G161 to E were 5′-CCATAGGATTTGAGAACATCTCACCACGC-3′ (forward) and 5′-GGGTGGTGAGATGTTCTCAAATCCTATGG-3′ (reverse), and 5′-CTCTAACAACTATTGAATTTGGTGACTACGTTGC-3′ (forward) and 5′-GCAACGTAGTCACCAAATTCAATAGTTGTTAGAG-3′ (reverse) for G268 to E. This mutant channel expresses well but carries no current when expressed in CHO cells.

