## Supplementals for "Coding of social novelty in the hippocampal CA2 region and its disruption and rescue in a mouse model of schizophrenia"

| Mean Firing Rate | Mean | SEM | ANOVA | WT vs Df(16) | Df(16) vs Spadin | WT vs Spadin |
|---|---|---|---|---|---|---|
| WT | 1.73 | 0.1322 | p=0.02 | p=0.02 | p=0.01 | p=0.57 |
| Df(16)A+/- | 1.3 | 0.1206 | | | | |
| Spadin | 1.87 | 0.2137 | | | | |

| Number of Fields | Mean | SEM | ANOVA | WT vs Df(16) | Df(16) vs Spadin | WT vs Spadin |
|---|---|---|---|---|---|---|
| WT | 2.8067 | 0.0778 | p<0.0001 | p<0.0001 | p<0.0001 | p=0.26 |
| Df(16)A+/- | 1.8923 | 0.0783 | | | | |
| Spadin | 3.0703 | 0.1115 | | | | |

| Field Size | Mean | SEM | ANOVA | WT vs Df(16) | Df(16) vs Spadin | WT vs Spadin |
|---|---|---|---|---|---|---|
| WT | 36.67 | .8583 | p=0.0009 | p=0.02 | p=0.02 | p=0.73 |
| Df(16)A+/- | 40.24 | 1.206 | | | | |
| Spadin | 36.16 | 1.258 | | | | |

| Selectivity | Mean | SEM | ANOVA | WT vs Df(16) | Df(16) vs Spadin | WT vs Spadin |
|---|---|---|---|---|---|---|
| WT | 10.61 | 0.9467 | p=0.0009 | p=0.02 | p=0.02 | p=0.80 |
| Df(16)A+/- | 13.96 | 1.044 | | | | |
| Spadin | 10.24 | 0.8526 | | | | |

| Spatial Info | Mean | SEM | ANOVA | WT vs Df(16) | Df(16) vs Spadin | WT vs Spadin |
|---|---|---|---|---|---|---|
| WT | .5453 | .0509 | p=0.01 | p=0.008 | p=0.03 | p=0.76 |
| Df(16)A+/- | .7365 | .0492 | | | | |
| Spadin | .5699 | .05173 | | | | |

**Supplemental Table 1. Spatial measures for the three CA2 experimental groups.**

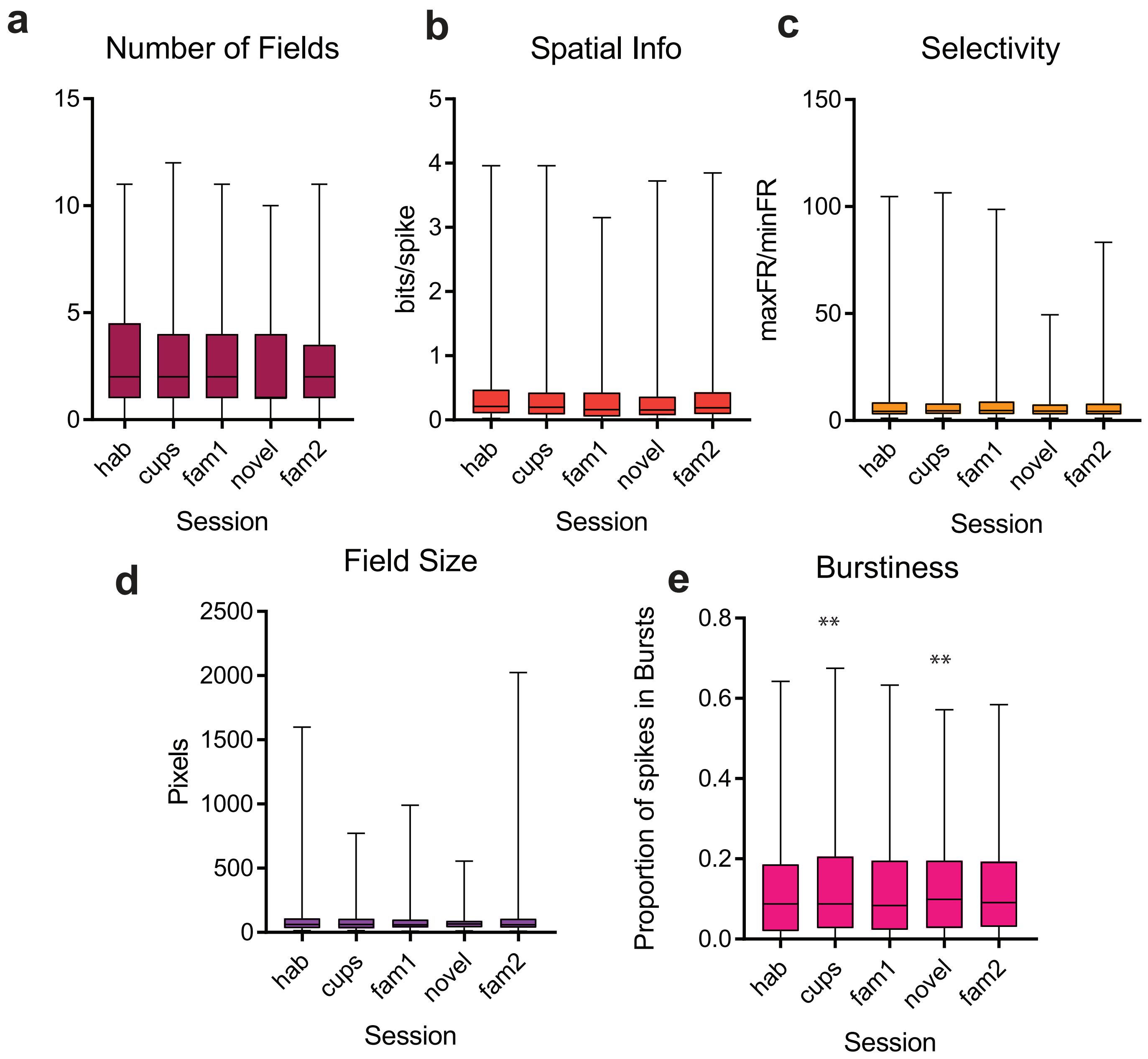


**Supplemental Figure 1. Single CA2 cell spatial firing properties during the different sessions of the three- chamber task.** (a-d) Single cell measures that did not differ during sessions (ANOVA $p>0.05$; $n=192$ CA2 neurons from 6 mice). (e) Burst index (number of spikes in bursts of at least three successive spikes with an interspike interval < 6 ms) was significantly higher in sessions with novel objects and the novel mouse ($p<0.01$, t-test post hoc to ANOVA).

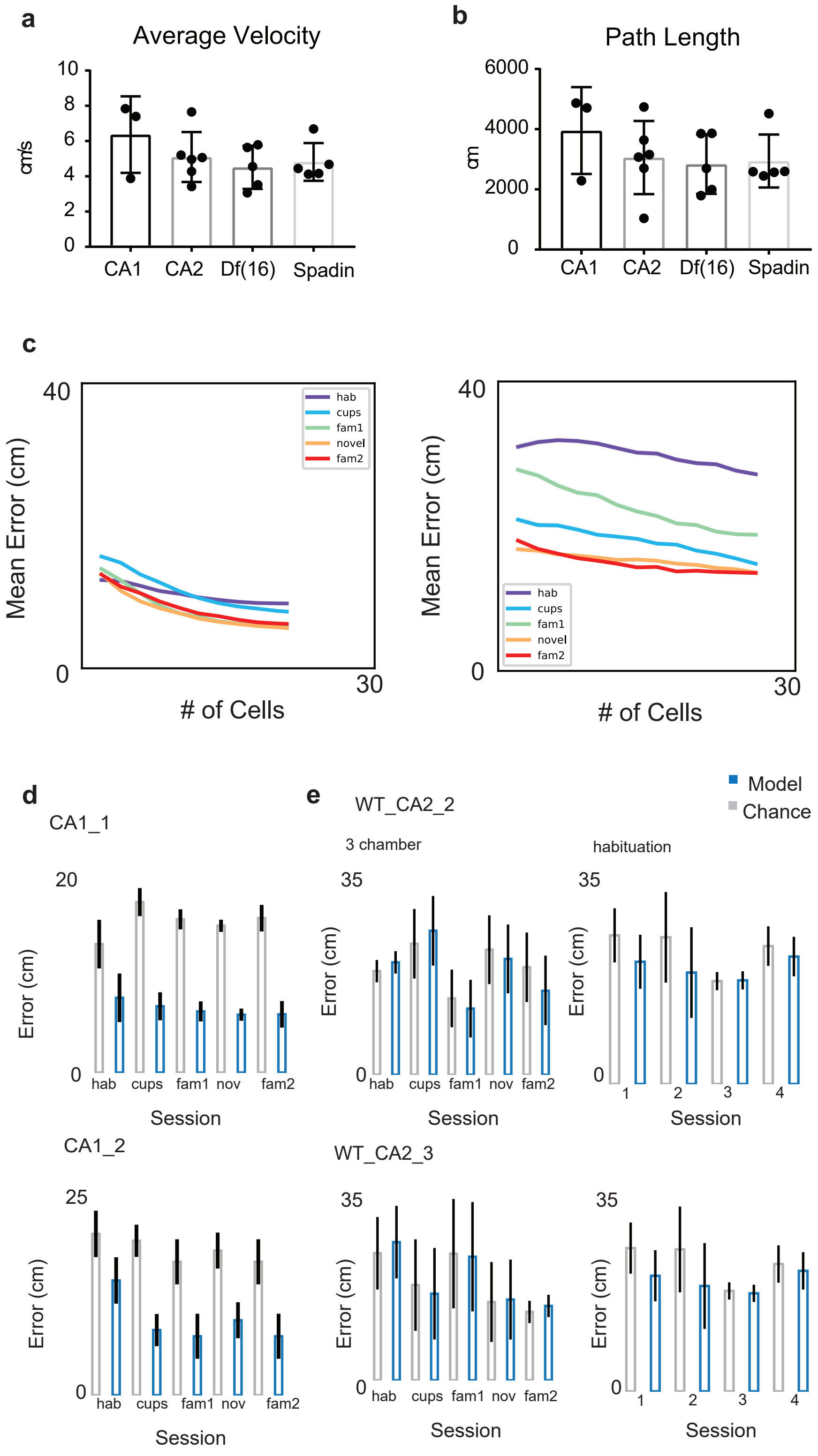


**Supplemental Figure 2. Mouse movement behavior and spatial decoding properties in various conditions**. There was no significant difference between (a) mean velocity (ANOVA, p=0.36), or (b) average path length (ANOVA, p= 0.71) in the different experimental groups of mice. (c) The ability to decode position increased as more CA1 (left) or CA2 cells (right) were added into the analysis. However, even with 30 cells CA2 spatial decoding accuracy was less than CA1 decoding accuracy with half as many cells. (d) CA1 population activity decoded position significantly better than chance in all sessions of the 3-chamber task. Data shown for two mice (CA1_1, n=21 neurons; CA1_2, n= 27 neurons). (e) CA2 population activity did not decode position better than chance in any 3-chamber task session (left graphs) or during the four ten-minute sessions of the habituation session (right graphs). Data shown for two mice (CA2_2, n= 25 neurons; CA2_3, n= 31 neurons).

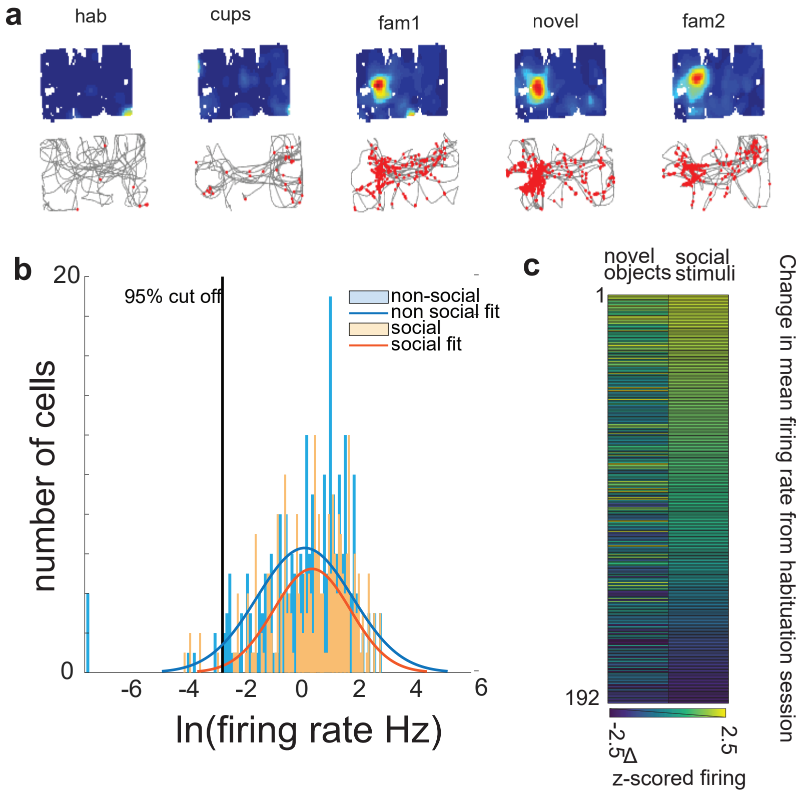


**Supplemental Figure 3. A subset of CA2 cells significantly increased their firing from the non-social to the social sessions.** (a) An example cell that began to fire in the social sessions. (b) Firing rate distributions for all cells in nonsocial (blue bars) and social (orange bars) sessions fit with Gaussian distributions. 6% of CA2 cells classified as silent in the non-social sessions, based on firing rates >2 SD below the median firing rate (<0.007 Hz), began to fire in the social sessions. (c) Change in z-scored firing rate from the empty arena session to the social sessions (right) and to the novel object session (left). The two firing rate vectors differed significantly (Wilcoxon rank-sum test, p=5.80 e-37).

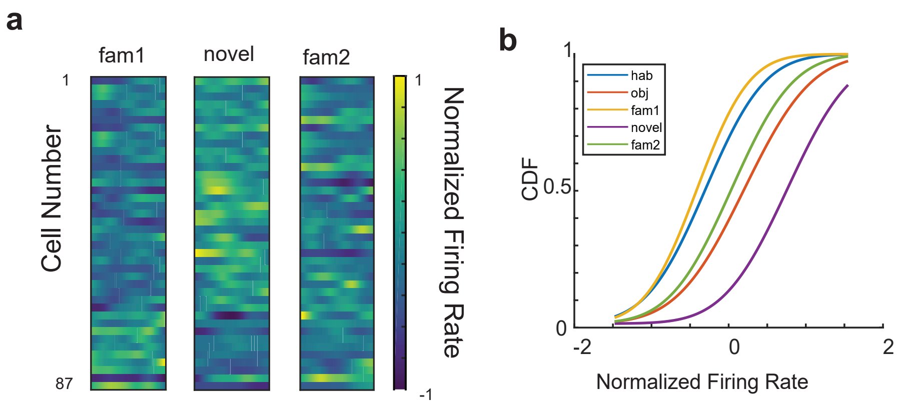


**Supplemental Figure 4. Responses of CA1 and CA2 neurons to novel stimuli.** (a) Color-coded z-scored firing rates in the interaction zone around the novel mouse and familiar mouse in sessions 3-5 for 87 CA1 neurons (n=3 mice). There was no significant difference in firing rates among sessions (ANOVA, p=0.17). (b) Cumulative distributions of z-scored firing rates for all 192 CA2 neurons (n= 6 mice) in the interaction zones in each session of the five 3-chamber task sessions. CA2 activity in the interaction zones was significantly different in the different sessions (Kruskal Wallis test; p <0.0001). CA2 firing rates in interaction zone around the novel object (wire cup cages) were significantly greater than in equivalent interaction zone in empty chamber (Mann Whitney, p<0.05). CA2 firing rates around the novel mouse were significantly higher than around the novel object (mean z-scored firing rates = 0.83 ± .06 and 0.19 ± 0.08, respectively; Mann Whitney U, p < 0.05).

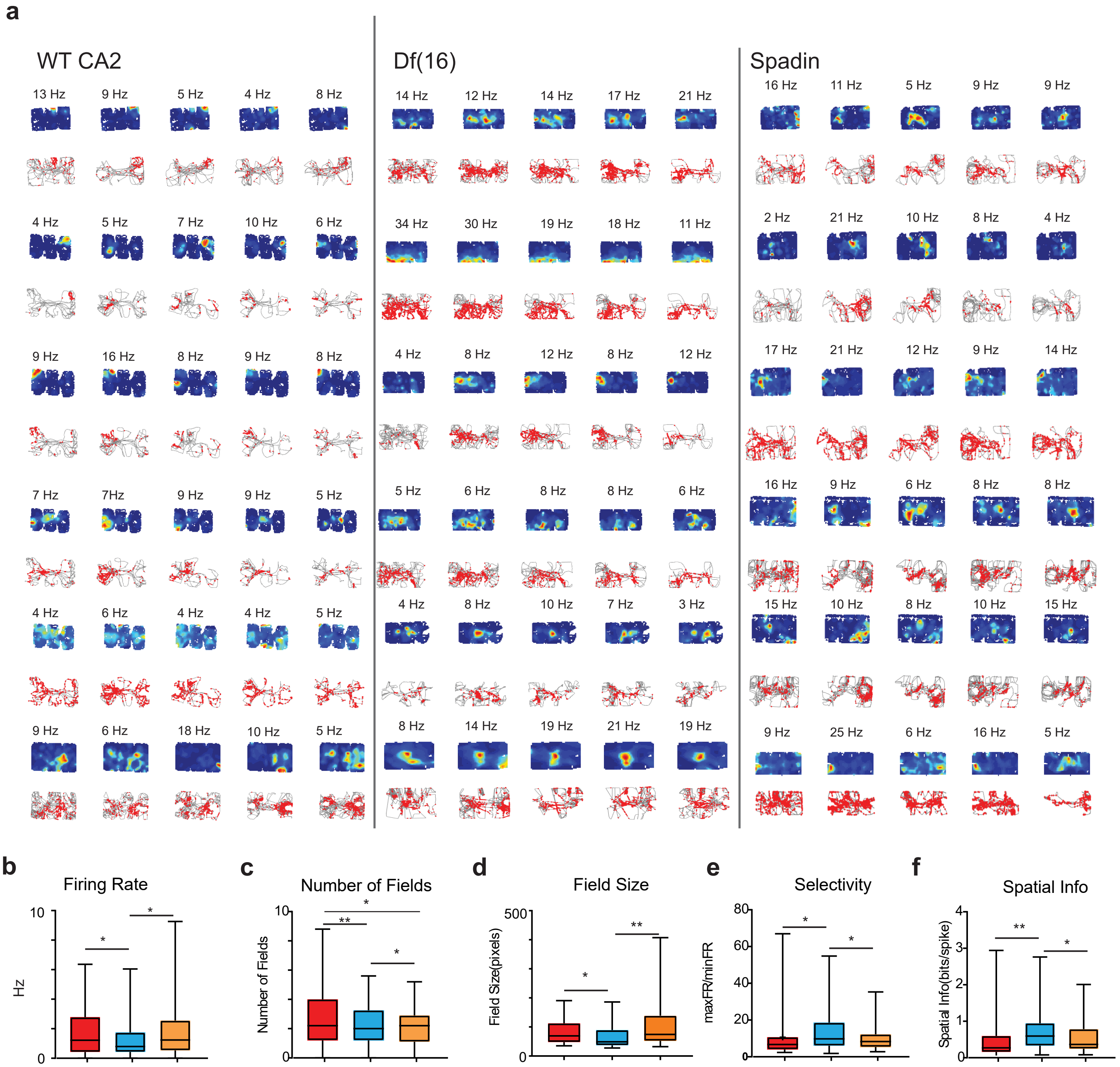


**Supplemental Figure 5. CA2 spatial firing in Df(16)A+/- mice and effect of spadin.** (a) Additional examples of CA2 neuron spatial firing from wild-type, Df(16)A+/-, and spadin-treated Df(16)A+/- animals. (b) CA2 neurons in Df(16)A+/- mice have a lower overall firing rate than wild-type mice (p=0.02, paired t-test) Spadin increased the firing rate in the Df(16)A+/- mice (p=0.04, paired t-test) (c) Compared to CA2 neurons in wild-type mice, CA2 neurons in Df(16)A+/- mice had: (c) fewer place fields per cell (p<0.0001, paired t-test); (d) smaller place fields (p=0.02); (e) place fields with higher spatial selectivity (p=0.02); and (f) higher spatial information content (p=0.008). CA2 neuron firing rates, place field size, selectivity and spatial information content in Df(16)A+/- mice treated with spadin were not significantly different from values in wild-type mice (p>0.05). Number of fields per cell in spadin-treated Df(16)A+/- mice was significantly greater than in untreated Df(16)A+/- animals (p=0.01) but significantly less than in wild-type mice (p=0.02).

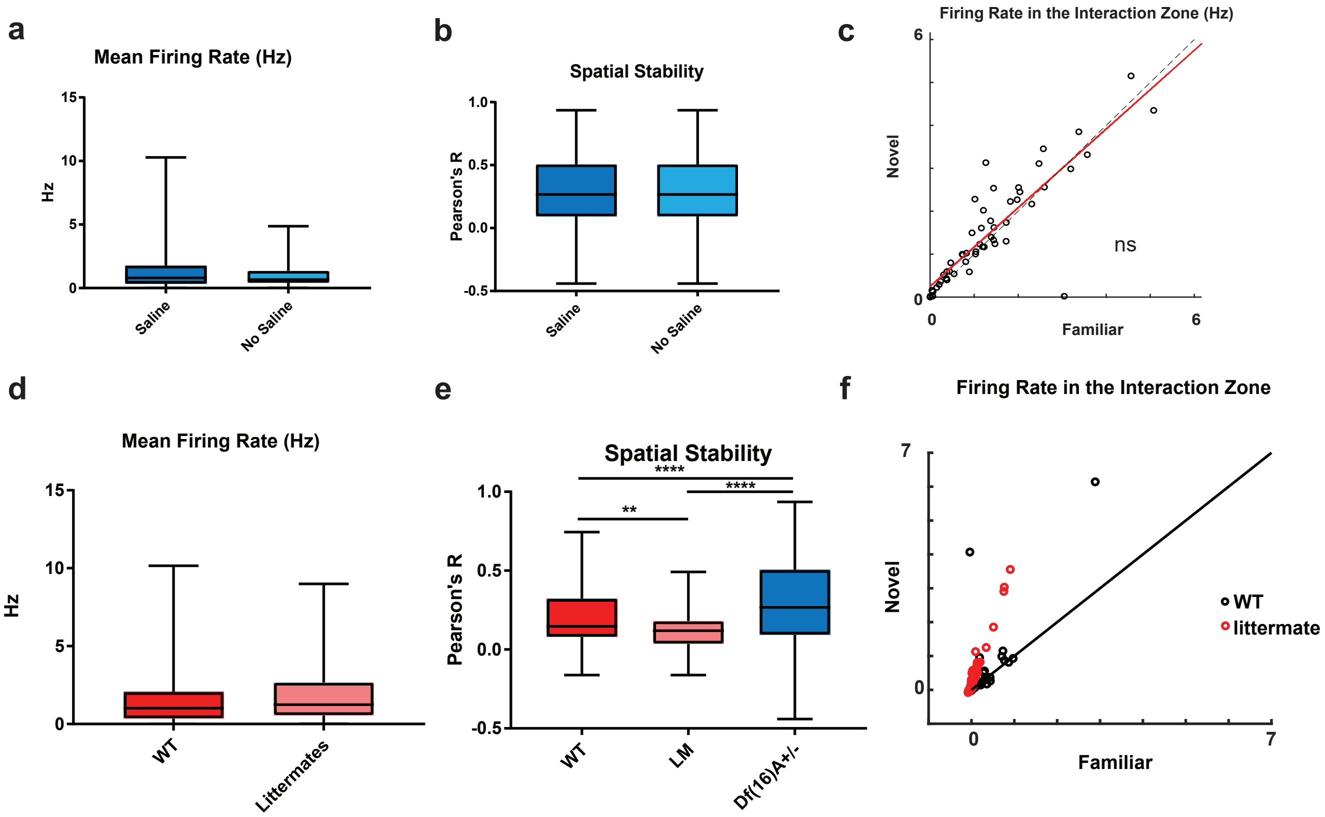


**Supplemental Figure 6. Control experiments for firing properties of CA2 neurons in Df(16)A+/- mice.** (a,b) Saline injection (control for spadin injection) in Df(16)A+/- mice did not alter CA2 neuron: (a) mean firing rate (paired t-test p=0.21) or (b) spatial stability in the five sessions of the three-chamber task (paired t-test, p=0.31). (c) CA2 neuron firing rate in saline-injected Df(16)A+/- mice did not differ in interaction zone around the novel compared to familiar mouse (p=0.7), similar to uninjected Df(16)A+/- mice (Figure 7a) but distinct from CA2 novel firing preference in wild-type (Figure 4c) and spadin-injected Df(16)A+/- mice (Figure 7d). (d-f) CA2 neuron firing properties in two groups of wild-type control mice used for comparison with Df(16)A+/- mice (all on identical C57Bl/6J backgrounds): wild-type littermates (n= 56 neurons from 2 mice) and wild-type non-littermates (n=136 neurons from 4 mice) . (d) There was no significant difference in mean firing rate between wild-type non-littermates (WT) and wild-type littermates (LM) (paired t-test, p= 0.19). (e) CA2 spatial stability was slightly but significantly lower in wild-type littermates compared to wild-type non-littermates (paired t-test with Bonferroni correction, p=0.002); spatial stability of both wild-type groups was significantly less than that of Df(16)A+/- mice (paired t-test with Bonferroni correction, p<0.0001 in both cases). (f) The two wild-type control groups showed a similar increase in firing around the novel animal compared to the familiar animal (p=0.45).

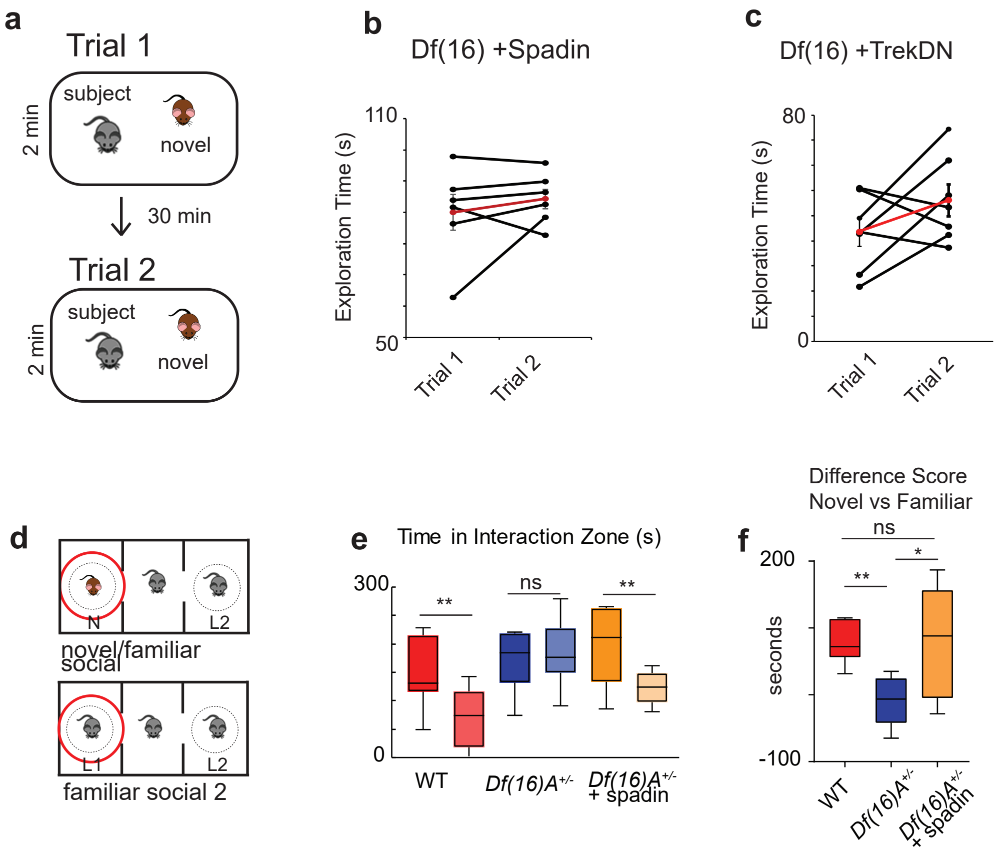


**Supplemental Figure 7. Effects of TREK-1 inhibition on social behavior**.(a) Control experiment for direct interaction test using a second novel mouse in trial 2 (instead of the same novel mouse used in trial 1). (b) There was no decrease in exploration of the second novel mouse when spadin was administered 30 min before trial 1 ($p=0.57$, paired t-test, n= 6 mice). (c) No decrease seen in exploration of second novel mouse in trial 2 in Df(16)A+/- mice expressing TREK-1 DN in CA2 ($p=0.34$, paired t-test, n= 6 mice). Black points and lines show individual animals. Red lines and points show means. Bars show SEM. (d,e,f) Comparison of social interaction times in three-chamber task for wild-type mice, Df(16)A+/- mice, and Df(16)A+/- mice injected with spadin (n=6, 5 and 5 mice, respectively). Time spent in interaction zone around cup containing the novel animal (novel session 4; bars with dark shades of color) compared to time spent in same interaction zone when the familiar mouse is present (averaged from familiar sessions 3 and 5; bars with light shades of color). Wild-type and spadin-treated Df(16)A+/- groups spent significantly less time exploring the familiar animal (paired t-test, $p<0.01$). Df(16)A+/- mice spent similar times exploring the familiar and novel animals ($p>0.05$; paired t-test). (f) Data from (e) plotted as difference scores. (ANOVA: WT versus Df(16)A+/- mice , $p<0.01$; Df(16)A+/- mice in absence versus presence of spadin, $p<0.05$).

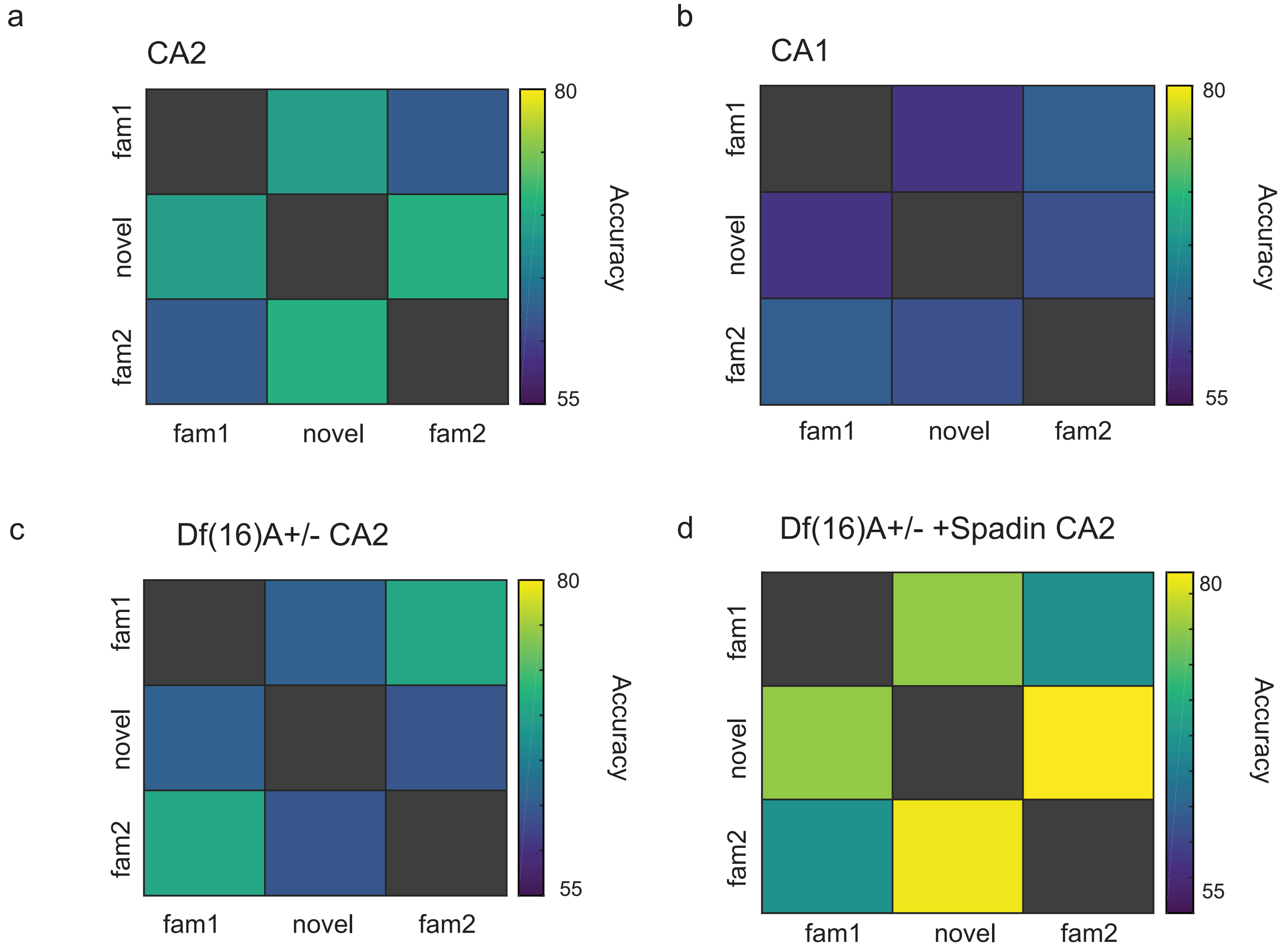


**Supplemental Figure 8. Color-coded plots of decoder accuracy**. Decoder was trained on neuronal firing in same interaction zone surrounding novel mouse and familiar mouse in sessions 3-5 (fam1, novel, fam2 sessions) of the three-chamber task. Each square shows accuracy in distinguishing indicating pairs of sessions. (a) Performance based on CA2 firing from wild-type mice (n= 192 neurons from 6 mice). (b) Performance based on CA1 firing from wild-type mice (n= 87 neurons from 3 mice). (c) Performance based on CA2 neuron firing from Df(16)A+/- mice (n= 128 neurons from 5 mice). (d) Performance based on CA2 neuron firing from Df(16)A+/- mice injection with spadin (n= 91 neurons from 5 mice).

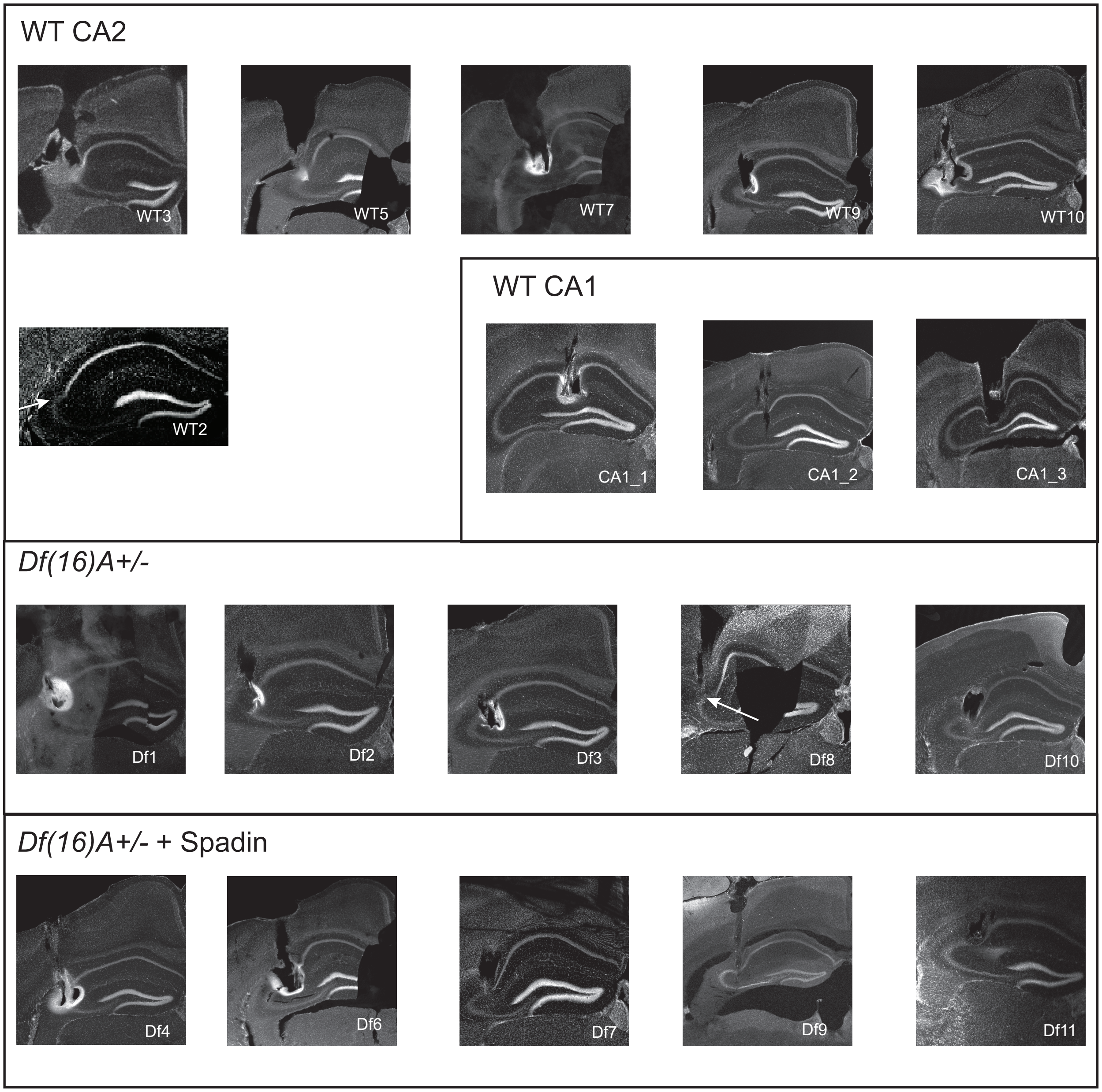


**Supplemental Figure 9. Tetrode tracks from all animals recorded in this study.**
